# Description of a novel extremophile green algae, *Chlamydomonas pacifica*, and its potential as a biotechnology host

**DOI:** 10.1101/2024.09.03.611117

**Authors:** João Vitor Dutra Molino, Aaron Oliver, Harish Sethuram, Kalisa Kang, Barbara Saucedo, Crisandra Jade Diaz, Abhishek Gupta, Lee Jong Jen, Yasin Torres-tiji, Nora Hidasi, Amr Badary, Hunter Jenkins, Francis J. Fields, Ryan Simkovsky, Stephen Mayfield

**Affiliations:** Department of Molecular Biology, School of Biological Sciences, University of California San Diego, San Diego, CA, United States; Center for Marine Biotechnology and Biomedicine, Scripps Institution of Oceanography, University of California San Diego, La Jolla, CA, United States; Infectious Disease and Microbiome Program, Broad Institute of MIT and Harvard, Cambridge, MA, USA; Microbial and Environmental Genomics Department, J. Craig Venter Institute, La Jolla, CA, USA; Colorado School of Mines, Golden, CO, USA; Department of Biology and Biotechnologies “Charles Darwin”, Sapienza University of Rome, Piazzale Aldo Moro, 5, 00185, Rome, Italy; Division of Pulmonary Diseases and Critical Care Medicine, University of California, Irvine, CA, United States; UCI Sleep Disorders Center, University of California, Irvine, 20350 Birch St., Newport Beach, CA, 92660, USA; Algenesis Inc., 1238 Sea Village Dr., Cardiff, CA, USA; California Center for Algae Biotechnology, University of California, San Diego, La Jolla, CA, United States; Sarawak Biodiversity Centre, KM20, Jalan Puncak Borneo, 93250 Kuching, Sarawak, Malaysia

**Keywords:** Chlorophyte, alkali tolerant, microalgal chassis, mating, stress tolerance (temperature, salinity), biotechnology, microalgae, new species, extremophile, bioplastic

## Abstract

We present the comprehensive characterization of a newly identified microalga, *Chlamydomonas pacifica*, originally isolated from a soil sample in San Diego, CA, USA. This species showcases remarkable biological versatility, including a broad pH range tolerance (6-11.5), high thermal tolerance (up to 42°C), and salinity resilience (up to 2% NaCl). Its amenability to genetic manipulation and sexual reproduction via mating, particularly between the two opposing strains CC-5697 & CC-5699, now publicly available through the Chlamydomonas Resource Center, underscores its potential as a biotechnological chassis. The biological assessment of *C. pacifica* revealed versatile metabolic capabilities, including diverse nitrogen assimilation capability, motility and phototaxis. Genomic and transcriptomic analyses identified 17,829 genes within a 121 Mb genome, featuring a GC content of 61%. The codon usage of *C. pacifica* closely mirrors that of *C. reinhardtii*, indicating a conserved genetic architecture that supports a trend in codon preference with minor variations. Phylogenetic analyses position *C. pacifica* within the core-Reinhardtinia clade yet distinct from known Volvocales species. The lipidomic data revealed an abundance of triacylglycerols (TAGs), promising for biofuel applications and lipids for health-related benefits. Our investigation lays the groundwork for exploiting *C. pacifica* in biotechnological applications, from biofuel generation to synthesizing biodegradable plastics, positioning it as a versatile host for future bioengineering endeavors.

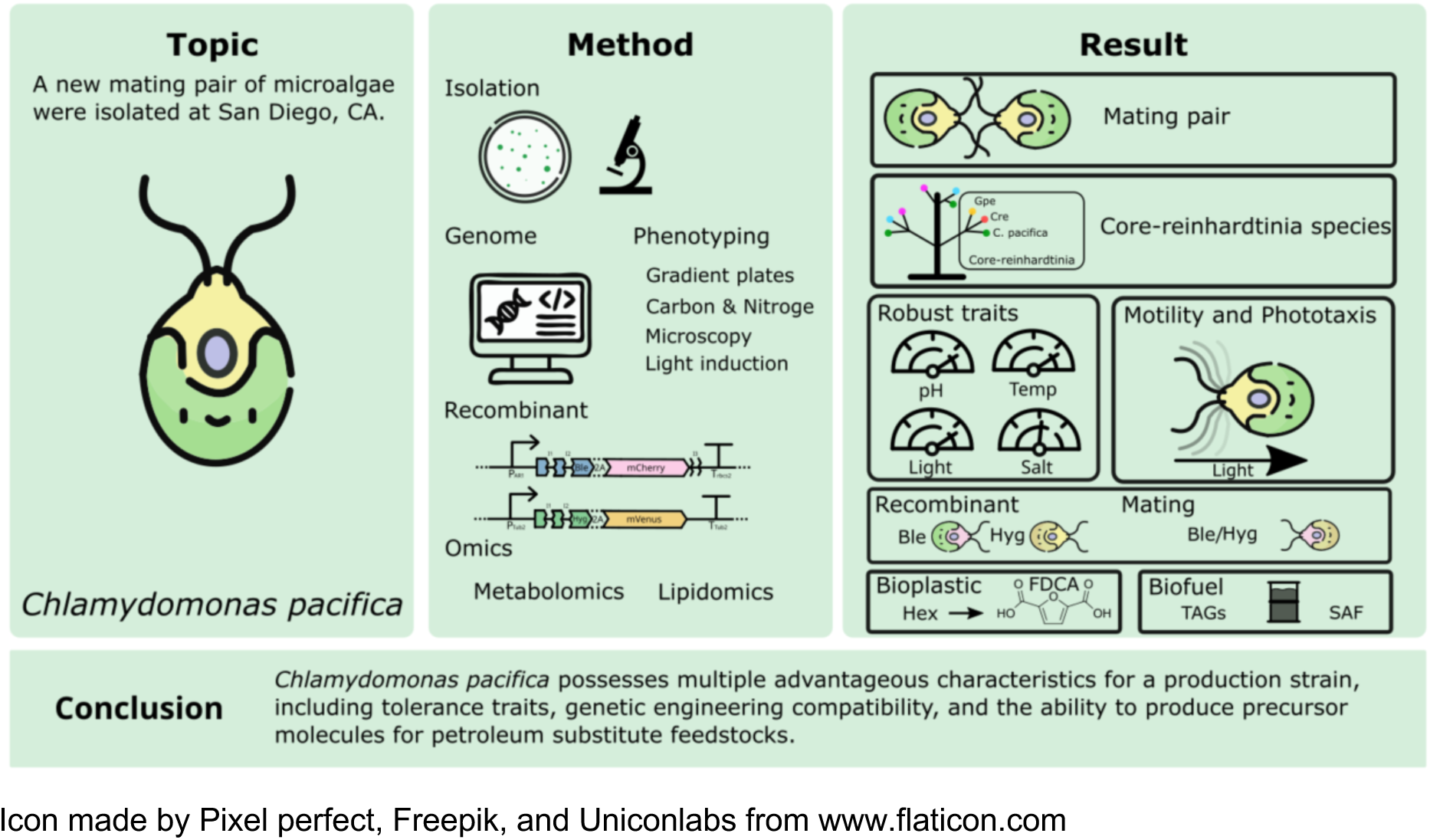
Graphical Abstract.

## Introduction

Green algae are generally aquatic plant-like organisms found in various environments, including freshwater, marine, and terrestrial habitats, and are the closest related species to land plants (Delwiche & Cooper, 2015). Their diversity is also reflected in the potential applications within biotechnology, where numerous species, including unicellular *Chlorella vulgaris*, *Dunaliela salina*, and *Chlamydomonas reinhardtii*, are currently employed (Kumar et al., 2020). Additionally, macroalgae such as *Ulva lactuca* and *Enteromorpha spp*., commonly used in East Asian cuisines, have gained some attention for their biotechnological potential (Ren et al., 2022). The global algae products market is estimated to be valued at USD 1.11 billion in 2024, reflecting a Compound Annual Growth Rate (CAGR) of 7.39% (TMR, 2016). This underscores the burgeoning significance of algae-based applications across various industries. However, the reliable cultivation and production of these diverse species requires specific growing techniques, with some species relying on specialized equipment, such as closed photobioreactors, to prevent contamination of the cultures (Novoveská et al., 2023). While algal systems are increasingly employed on large scales, their natural diversity is significantly broader. For instance, the genus Chlamydomonas contains more than 450 species reported (Milo et al., 2010), yet only a handful of species are used commercially (Tran et al., 2019). This is due to the suitability of specific species for producing desired products, as applications like nutraceuticals or functional foods that can accommodate higher production costs (Diaz et al., 2023; Ejike et al., 2023; Udayan et al., 2023; Vishwakarma et al., 2023), while others require lower costs, such as wastewater treatment, biofuel, bioplastic production, and animal feed (Abdelfattah et al., 2023).

One notable advantage of algae cultivation is its ability to thrive in diverse environments, including non-arable lands and wastewater treatment facilities, thereby avoiding competition with food crop cultivation (Ahmad & Ashraf, 2023). Another advantage this crop holds over traditional terrestrial plants is its remarkable ability to thrive in various water sources, including seawater and brackish water, the latter being a largely untapped resource where conventional crops typically fail to flourish. For example, in the United States alone, there is more than 800 times the amount of water currently used yearly in the form of brackish water bodies (Stanton et al., 2017). The location of the main basins also coincides with regions characterized by non-arable land, such as Arizona, New Mexico, Utah, Nevada, California, and parts of Colorado— all benefiting from high sunlight incidence, making them ideal for algae cultivation. To achieve industrial-scale production, it is essential to select strains that exhibit consistent growth and can thrive in challenging conditions, including high light incidences, elevated temperatures, and brackish water. These regions are often underutilized for traditional land use or pastureland and represent untapped potential for algae cultivation. The ability of a strain to flourish in such environments not only expands the geographic scope for algae cultivation but also provides a strategic advantage in resource utilization.

The cost associated with cultivating a species is inherently tied to its traits and growth requirements. While *C. reinhardtii* can be grown on an industrial scale, its production is typically accomplished within closed bioreactors using an organic carbon source such as acetic acid (Torres-Tiji et al., 2022). These closed systems provide conditions conducive to growth that are otherwise unsuitable for open systems. Therefore, the selection of algae strains plays a crucial role in the choice and feasibility of production systems. Growing in extreme conditions, which prevents the risk of contamination, favors using cheaper and more scalable systems like the high-salinity green algae *D. salina*, capable of thriving in 85 g/L of salt (B. Zhang et al., 2023). This adaptability simplifies management efforts and helps lower production costs (Varshney et al., 2015a). Extremophiles are found in various extreme environments, including hot springs, ice caps, high salt (*Dunaliella* sp.), and acidic mines and rivers (*C. acidophila*) or soda lakes (*Spirulina platensis*) (Brock, 1975; Luís et al., 2022; Melack, 1979; Oren, 2014; Remias et al., 2005; Selvaratnam et al., 2014). Particular adaptations allow them to survive in harsh conditions, making them interesting organisms for bioprospecting. On one hand, extremophiles can serve as sources of valuable tools, such as Taq polymerase; on the other hand, their inherent ability to grow in extreme conditions simplifies culturing management. For instance, maintaining them in high-salt conditions can help prevent contamination and protect against predators, addressing a significant obstacle faced in the cultivation process (McGowen, 2019; Varshney et al., 2015b).

This study presents *Chlamydomonas pacifica*, a newly isolated extremophile green alga capable of thriving under high pH, elevated temperatures, intense light irradiance, and brackish water salinity. We conducted a comprehensive multi-omics characterization to elucidate its genetic and metabolic capabilities, including genome sequencing, transcriptomics, and lipidomics. These analyses revealed a diverse lipid profile with potential applications in biofuels and health-related industries. In addition to multi-omics approaches, we performed extensive phenotypic evaluations to characterize its tolerance to pH, salinity, temperature, and light intensity, as well as its morphology, motility, phototaxis, and mating behavior. Two mating types capable of sexual reproduction were isolated and genetically engineered with distinct expression vectors. We also evaluated the strains’ ability to assimilate diverse nitrogen and carbon sources. Collectively, these findings establish a foundation for developing *C. pacifica* as a robust platform for biotechnological applications, contributing to a sustainable bioeconomy.

## Results and Discussion

### Genome features

We assembled and annotated the genome of *Chlamydomonas pacifica* to investigate its unique features and gain deeper insights into its biology. *C. pacifica* exhibits a genome size of 121 Mbp and a lower GC content (61%) compared to *C. reinhardtii*. To assess the completeness of the annotated *C. pacifica* genome, Benchmarking Universal Single-Copy Orthologs (BUSCO), a tool that evaluates the presence of a highly conserved set of single-copy genes in most Chlorophyta lineages, was employed. The BUSCO analysis revealed 91.3% completeness, indicating minimal gene loss and the comprehensiveness of the genome annotation. We identified 17,829 protein-coding genes in *Chlamydomonas pacifica*, a number comparable to its well-studied counterpart, *C. reinhardtii* (16,795 genes in CC-503 v6.1) (Craig et al., 2023). Using the annotated genes, we generated a codon usage table for *C. pacifica* (**Supplementary Table 1**) and compared it with *C. reinhardtii*. The analysis revealed a linear association, with a regression coefficient of 0.92 and high correlation (R² = 0.95) in codon usage patterns between the two species (**Supplementary** Figure 1). Notable differences were observed in the stop codons TAA and TGA, as well as the lysine codons AAG and AAA. However, the majority of codon frequencies (57 out of 64) differed by less than 10%, highlighting a high degree of similarity in codon usage between the two species. This similarity suggests that DNA sequences optimized for *C. reinhardtii* will likely be efficiently translated into *C. pacifica*. The shared codon bias facilitates genetic engineering by allowing the use of the extensive repertoire of expression vectors already developed for *C. reinhardtii*, a capability we experimentally validated (**see Genetic Engineering section**).

We then constructed a phylogenetic tree (**Figure 1**) using 1,519 chlorophyta genes and 42 available algae species to place *Chlamydomonas pacifica* on the Core-reinhardtinia, of which *C. reinhardtii* is a distinguished member (Craig et al., 2023). The closest relative to *Chlamydomonas pacifica* is *Chlamydomonas sp. 3222*, a strain isolated in the United Arab Emirates (Nelson et al., 2019). This strain, available in the Chlamydomonas Resource Center under the index CC-5164, was obtained for comparative analyses with *C. pacifica* to explore their physiological similarities further.

**Figure 1:**
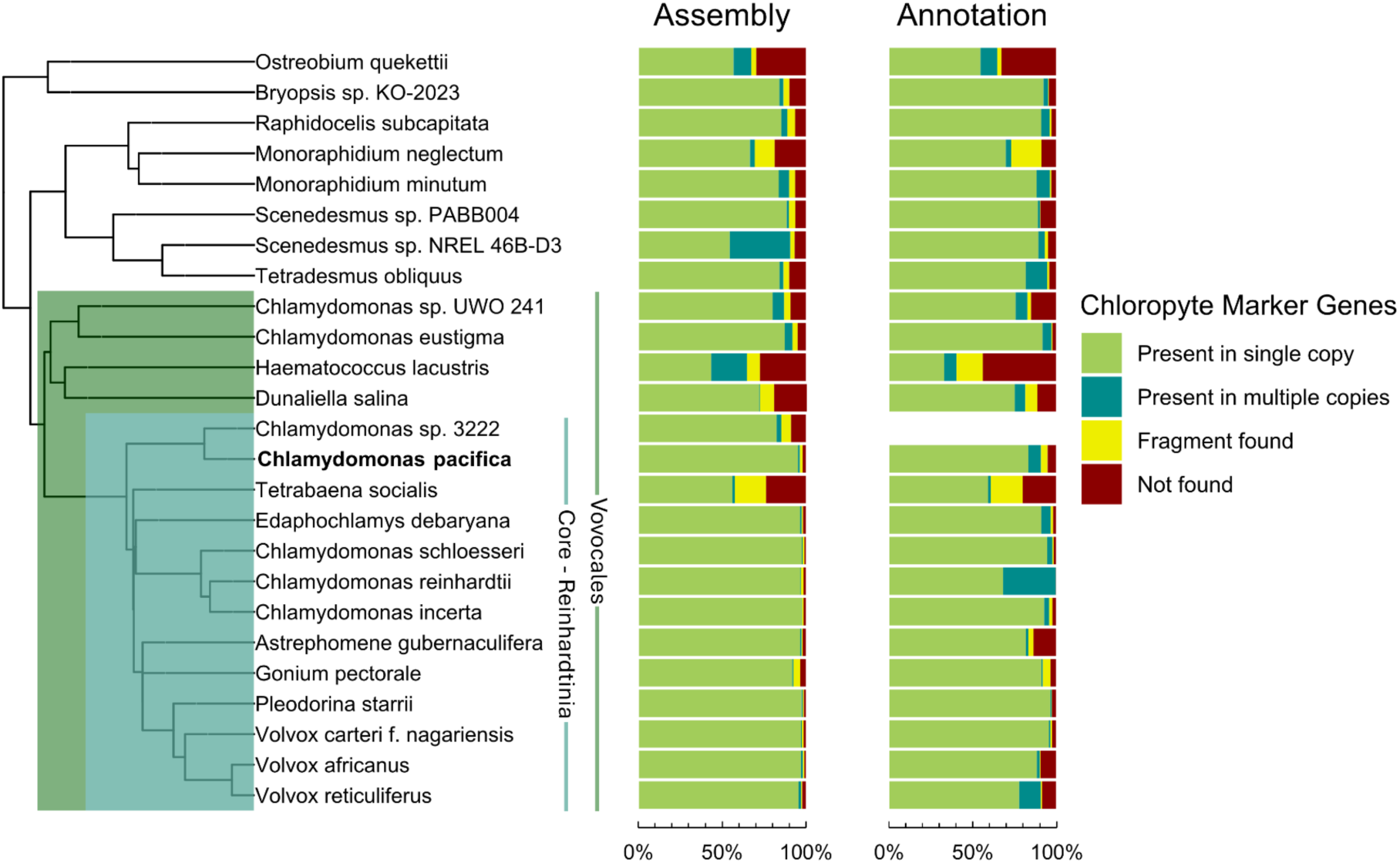
Phylogenetic distribution of assembled chlorophyte genomes based on average amino acid identity. Completeness bar graphs adjacent to the phylogenetic tree display BUSCO scores for the genome assembly and gene models, if publicly available. *C. pacifica* completeness of assembled and annotated chlorophyte genomes is consistent with other members of the Vovocales (Seppey et al., 2019).

The study also assessed the occurrence of genes involved in mating processes, revealing homologies between *C. reinhardtii* and *C. pacifica* in several mating-related genes (**Supplementary Table 2**) (Ferris et al., 2002a). These include genes unique to MT-gametes (mtd1, mid), a MID-dependent activation protein (GSM1), the sexual agglutination gene (SAG1), sexual adhesion gene (SAD1), the HAP2 gene responsible for membrane pore formation during mating, the nuclear fusion protein GEX1, the gametogenesis-active matrix metalloprotease gene Mmp1, the mating type plus-specific protein GSP1 active in zygotes, and the gamete fusion protein Fus1 (Kubo et al., 2001; Lin & Goodenough, 2007; Ning et al., 2013; Pinello & Clark, 2022). The sequence similarity among these genes varied significantly, with identity ranging from 4.6% to 37.1%, indicating diverse evolutionary rates (**Supplementary Table 2**). The presence of mating genes does not necessarily equate to mating capability, at least under laboratory conditions, as has been observed in other species (Blanc et al., 2010). Nevertheless, we successfully validated the mating capacity of the isolated *C. pacifica* strains through experimental assays (**See Mating section**).

The assembly and annotation of the *C. pacifica* genome provide critical insights into its biology and biotechnological potential. The genome facilitated the identification of its closest relative, *Chlamydomonas sp. 3222*, revealed a codon usage bias similar to *C. reinhardtii* and uncovered key mating-related genes. These features enhance the feasibility of genetic engineering and highlight *C. pacifica*’s potential as a valuable platform for biotechnology applications.

### Physiological and Environmental Tolerance Characterization

The four *Chlamydomonas* strains were comprehensively characterized, emphasizing their nutrient assimilation capabilities and environmental resilience. The analysis included *C. reinhardtii* as the well-studied reference species within the Core-reinhardtinia clade, *C. sp. 3222* as the closest related species (**see Phylogenetic tree, Figure 1**), and the two mating strains of *C. pacifica*, 402 and 403. This comprehensive assessment aims to delineate the physiological and ecological niches of these strains, with a particular focus on understanding the similarities and differences between the closely related *C. sp. 3222* and *C. pacifica* strains, thereby shedding light on their potential for adaptive strategies and applications in biotechnological settings.

#### Carbon and Nitrogen Assimilation

We investigated the growth of the four strains under distinct regimes. These conditions were designed to assess their photosynthetic and heterotrophic metabolic capacities. Growth on nitrogen-source plates under continuous light (80 µmol photons m⁻² s⁻¹) emphasized a predominantly photosynthetic mode (**Figure 2)** while evaluating the strains’ ability to assimilate various nitrogen sources, specifically NH₄⁺, NO₃⁻, and urea. Conversely, carbon-source plates were incubated under low light (approximately 8 µmol photons m⁻² s⁻¹), minimizing photosynthetic activity to assess the strains’ heterotrophic growth capacity (**Figure 3**) using acetate, glycerol, glucose, and sucrose as potential carbon sources. This approach provided a nuanced understanding of their metabolic flexibility and adaptation to varying energy sources.

**Figure 2.**
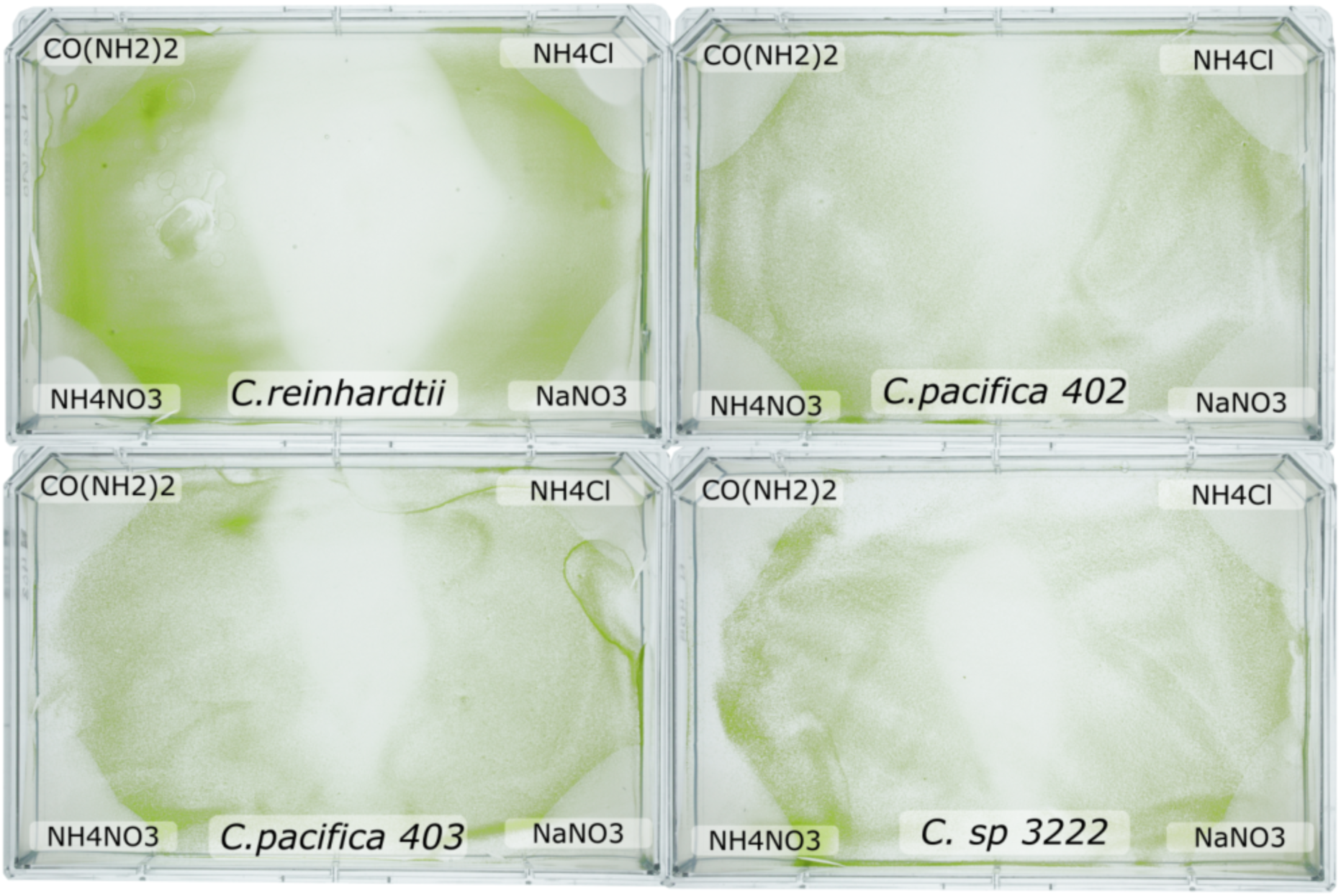
Comparative Growth Responses of *Chlamydomonas* Species to Different Nitrogen Sources. Petri dishes exhibit the growth of three *Chlamydomonas* strains: *C. reinhardtii*, *C. pacifica* strains 402 and 403, and *C. sp.* strain 3222, cultured in HSM without nitrogen media. The corners were supplemented with nitrogen compounds at 9.34 mM of nitrogen atoms. From left to right, the nitrogen sources provided are urea (CO(NH2)2), ammonium chloride (NH4Cl), sodium nitrate (NaNO3), and ammonium nitrate (NH4NO3). The center portion serves as a negative control.

**Figure 3.**
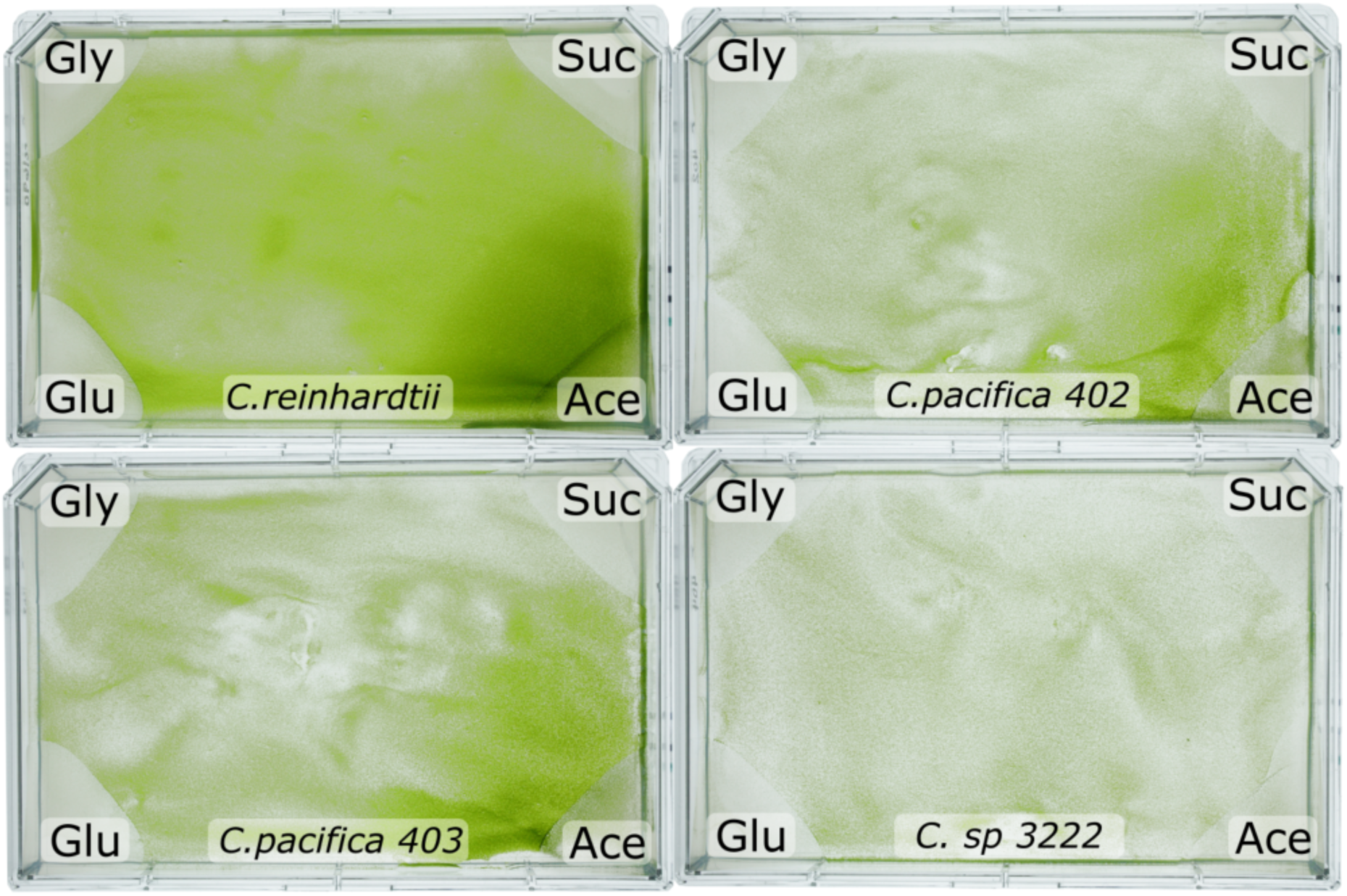
Differential Growth of *Chlamydomonas* Strains on Various Carbon Sources. The petri dishes demonstrate the growth patterns of *C. reinhardtii*, *C. pacifica* strains 402 and 403, and *C. sp.* strain 3222 when cultured on HSM media supplemented with distinct carbon sources. The carbon compounds tested include acetate (Ace), sucrose (Suc), glycerol (Gly), and glucose (Glu), arranged clockwise from the top left corner of each dish. The variation in green pigmentation intensity across the media indicates the efficiency of carbon utilization by each algal strain. Plates were incubated at room light to reduce photosynthetic growth.

Under continuous illumination, all strains demonstrated the capacity to assimilate the provided nitrogen sources (**Figure 2**), consistent with established research indicating efficient integration of such sources by Chlamydomonas species (Fernández et al., 2009). This versatility in nitrogen sources bears significance for large-scale algal culture systems, where choosing a nitrogen source can profoundly impact biotechnological processes’ sustainability and economic viability (Arumugam et al., 2013).

Conversely, the carbon-source assessment revealed that in the vicinity of the acetate, more growth was observed for *C. pacifica* strains, indicative of enhanced growth surpassing that attributable solely to photosynthesis (**Figure 3**). Acetate assimilation in these strains, potentially occurs via acetate activation and subsequent entry into the Tricarboxylic Acid Cycle (TCA) cycle. This metabolic pathway has been previously described in microalgae (Bogaert et al., 2019), and is a common trait in microalgae (Wolf et al., 2015). On the other hand, the lack of such intense growth near other carbon sources (sucrose, glycerol, and glucose) implies ineffective assimilation under the given conditions, with growth primarily reliant on photosynthesis, as evidenced by similar growth to the plate center (photosynthetic region). The *C. pacifica* strains demonstrated an ability to utilize acetate for growth, an attribute relevant to bioreactor systems where light is limited, and the cells can be grown heterotrophically to reach densities as high as 40g/L for *C. reinhardtii* (Torres-Tiji et al., 2022).

#### pH Profile

The pH tolerance of *Chlamydomonas* strains was assessed through a gradient plate assay (**Figure 4**), revealing significant variations in the capacity of these strains to thrive across different pH levels. The mesophilic nature of *C. reinhardtii* was observed to have optimal growth within a moderate pH range, aligning with previous characterizations of its preferred neutral environments (Harris et al., 2009b). This species’ adherence to mesophilic growth conditions highlights its evolutionary tuning to stable pH environments, a trait that could have implications for its ecological distribution and suitability for controlled bioprocesses. In contrast, the *C. pacifica* strains 402 and 403, alongside *C. sp.* strain 3222, exhibited a more extensive pH tolerance, thriving in conditions extending from pH 6 to greater than pH 10. The broader pH adaptability of these strains, especially when considered in conjunction with their phylogenetic proximity, underscores a potentially shared evolutionary capacity for environmental resilience. High pH tolerance could be leveraged in open pond cultures to protect the process from pests, a factor allowing commercial production of *Arthorspira platensis* (Spirulina) (Molina-Grima et al., 2022).

**Figure 4:**
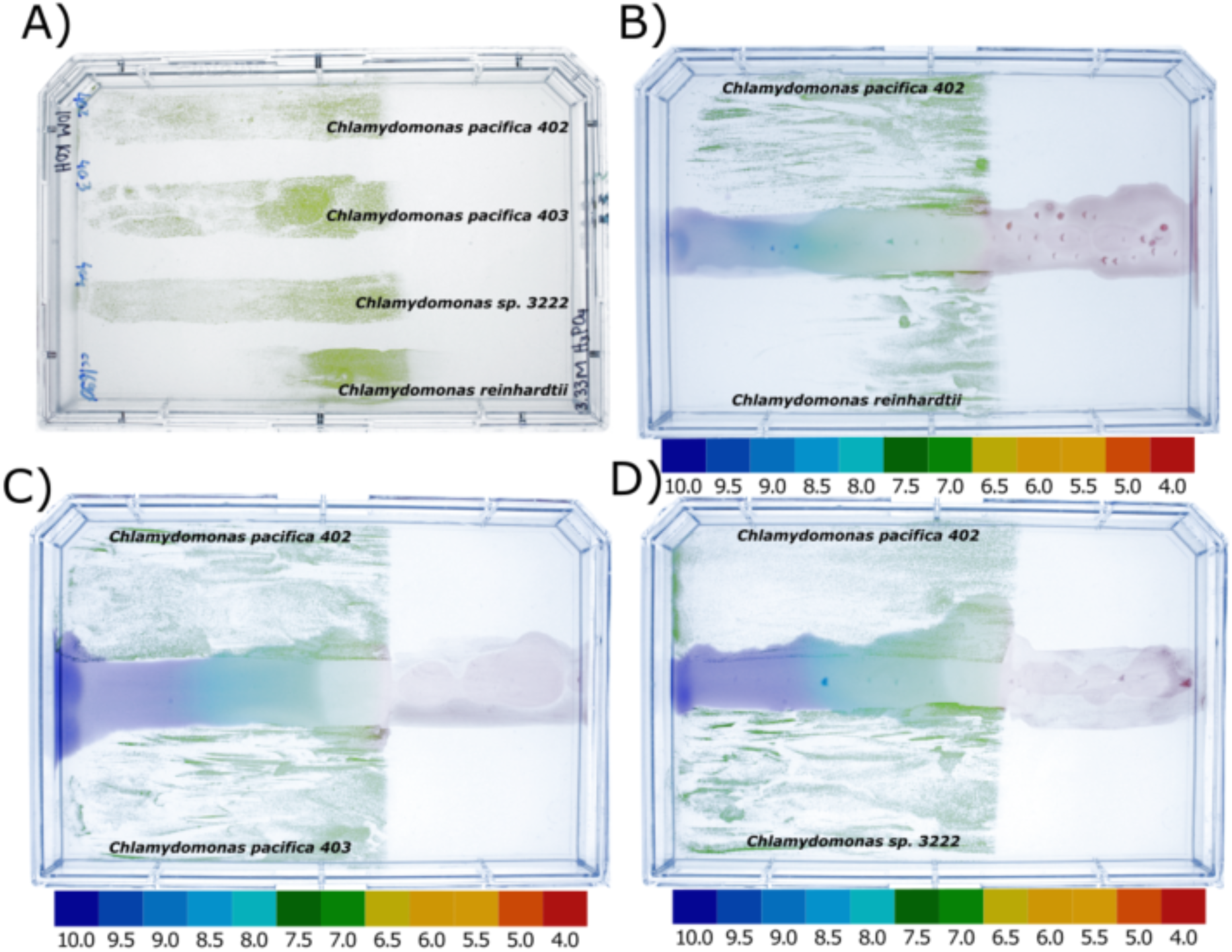
pH Gradient-Based Growth Profiling of *Chlamydomonas* Strains. The growth patterns of *C. reinhardtii*, *C. pacifica* strains 402 and 403, and *Chlamydomonas sp*. strain 3222 are demonstrated across a pH gradient. **A**) Growth of *C. pacifica* strains 402 and 403, *C. sp.* strain 3222, and *C. reinhardtii* across a pH gradient. **B**) *C. pacifica* strain 402 and *C. reinhardtii* growth response to the pH gradient. **C**) Growth of *C. pacifica* strains 402 and 403 across a pH gradient. **D**) *C. pacifica* strain 402 growth with C. sp. strain 3222 across a pH gradient. The gradient was established by applying 60 µL of 3.3M H3PO4 to the right side and 120 µL of 10 M KOH to the left side of each agar plate. A universal pH indicator (Fisher Chemical™ Universal pH Indicator System) was used to visualize the pH levels, with color changes indicating variations in pH across the plate. The differential growth observed reflects the varying capacities of the strains to tolerate and thrive under different pH conditions, providing insights into their adaptability and potential for cultivation in diverse environmental settings.

#### Salt Profile

To investigate the ability of the *Chlamydomonas* strains to withstand osmotic stress, we cultured them in HSM with varying concentrations of NaCl, which served to mimic different osmotic pressures found in water ecosystems (**Figure 5**). The strains *C. pacifica* 402 and 403 were found to sustain growth at salinity levels as high as 300 mM NaCl in addition to the media, with the 403 strain showing slightly better growth under these conditions. Osmotolerance was further assessed using NaCl gradient plates, where *C. pacifica* strains 402, 403, and *C. sp.* strain 3222 demonstrated comparable and significantly greater salt tolerance than *C. reinhardtii*, which exhibited growth over a narrower salinity range. The gradient plate experiments revealed that these three strains, including *C. sp.* strain 3222, possess similar capacities to thrive in brackish salinity conditions. For context, brackish groundwater in the United States typically contains 14.1 g/L of dissolved solids (Honarparvar et al., 2021). These findings point to a significant osmoregulatory ability in strains 402 and 403, as well as *C. sp.* strain 3222. Halotolerance is a critical trait for bioprocesses that rely on saline water sources, such as those associated with coastal or estuarine aquaculture systems. Additionally, it enables the utilization of brackish water sources in inland regions worldwide, which are typically unsuitable for conventional agricultural practices.

**Figure 5:**
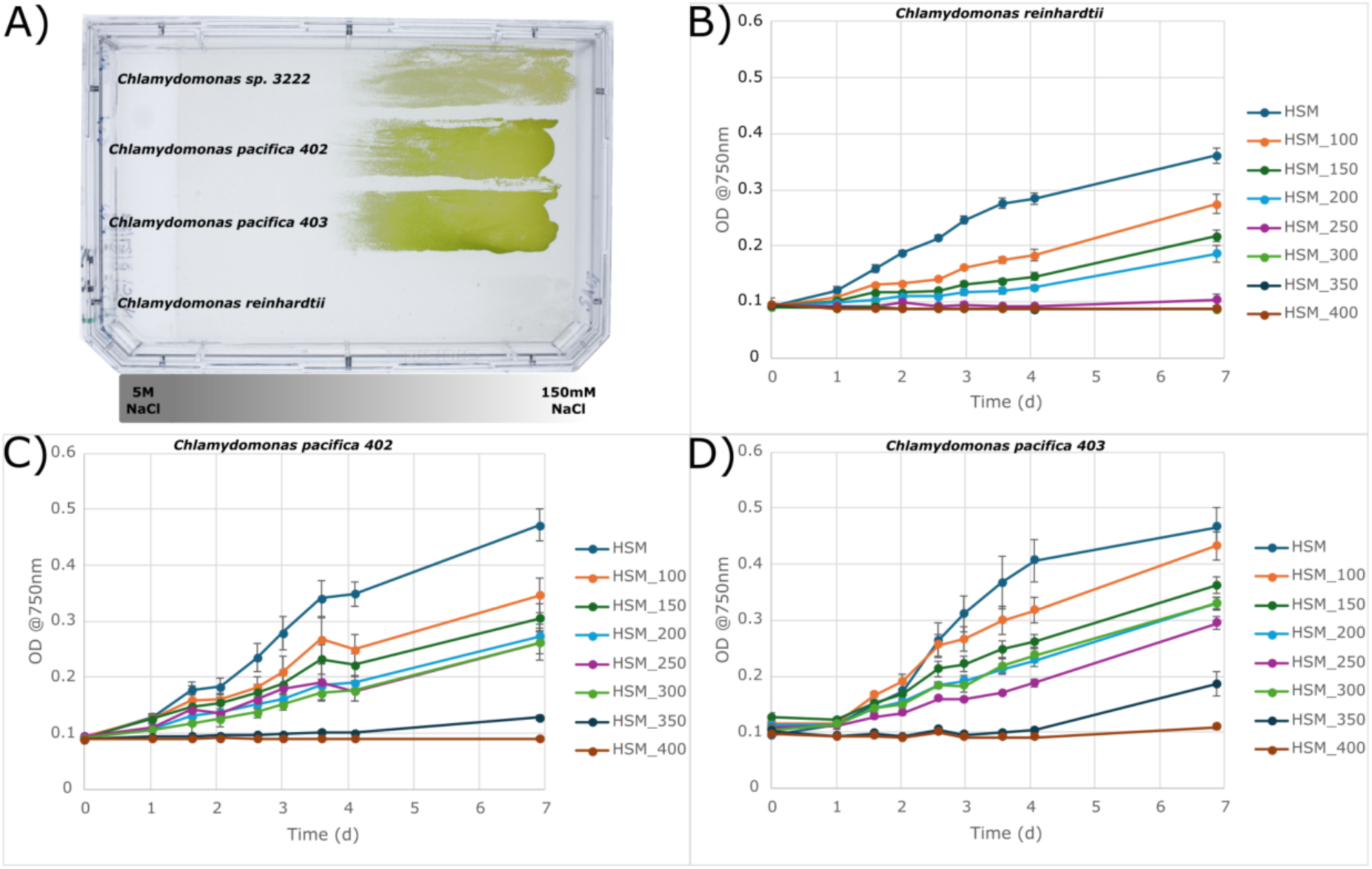
Salinity Tolerance of Chlamydomonas Strains. **A**) NaCl gradient plate delineating the comparative growth of *C. reinhardtii*, *C. pacifica* strains 402 and 403, and *C. sp.* strain 3222, with the left part containing high salinity (5 M NaCl) and the right side moderate salinity (150 mM NaCl). **B-D**) Growth curves over seven days for *C. reinhardtii* (**B**), *C. pacifica* 402 (**C**), and *C. pacifica* 403 (**D**) in high salt media (HSM) with incremental NaCl additions ranging from 0 to an additional 400 mM. The data illustrate the relative optical density (OD) growth at 750 nm.

#### Temperature Profile

We conducted a thermal tolerance assay on the Chlamydomonas strains to provide critical insights into their potential for cultivation in global regions with high-temperature variability, especially semi-arid areas where competition with agriculture is minimal. In these regions, temperatures frequently exceed 40°C (Winckelmann et al., 2015), necessitating robust thermotolerant strains for successful algal cultivation. *C. pacifica* strains 402 and 403, exhibited considerable resilience to elevated temperatures, withstanding up to 41.9°C (**Figure 6**), while *C. sp.* 3222 reached a maximum of 40.2°C. This degree of thermotolerance surpasses that of *C. reinhardtii* CC1690, which preferred cooler conditions (37.9°C), typical of its mesophilic nature. The ability to thrive at higher temperatures is a desirable trait for algae intended for biofuel production since it opens the possibility of exploiting solar energy in warm, arid regions that may not be suitable for traditional agriculture (Benemann, 2013; Tiwari et al., 2019). The high-temperature tolerance observed in *C. pacifica* strains and *C. sp.* strain 3222 suggests their suitability for deployment in these environments, where daytime temperatures can limit the productivity of less tolerant species. This advantage could enable continuous, year-round cultivation, contributing to the feasibility of commercial-scale algal biofuel projects in semi-arid and arid regions worldwide.

**Figure 6:**
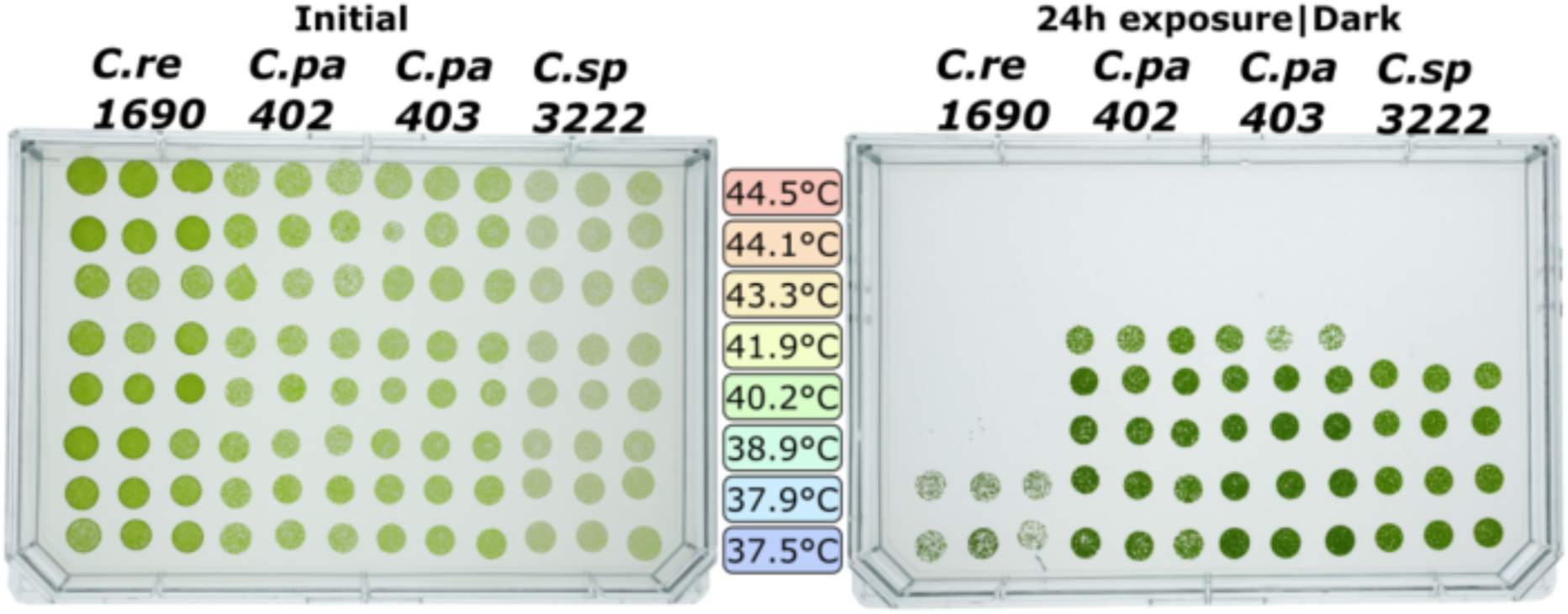
Thermal Tolerance of *Chlamydomonas* Strains in HSM Acetate Media. The left panel shows the initial state of *C. reinhardtii* CC1690, *C. pacifica* strains 402 and 403, and the phylogenetically related *C. sp.* strain 3222 before temperature treatment. The right panel displays the same strains post-24-hour exposure to a temperature gradient ranging from 37.5°C to 44.5°C, in ascending order from bottom to top, incubated in the dark. The cultures were placed in 100 µL aliquots within a 96-well PCR plate, which was then sealed to ensure containment. Triplicates for each strain and temperature combination were used to ensure experimental reliability. The pattern of growth post-exposure indicates the varying degrees of thermotolerance among the strains, with *C. pacifica* strains and C. sp. strain 3222 exhibiting greater resilience to higher temperatures compared to *C. reinhardtii* CC1690.

#### Light Tolerance Profile

The investigation into the light tolerance of *Chlamydomonas* strains has elucidated their potential for sustainable growth in high-irradiance environments characteristic of semi-arid regions, where high sunlight intensity can be a limiting factor for photosynthetic organisms. In these geographical areas, daily light intensities can frequently surpass 2000 µE/m²·s, demanding high resilience to photic stress for algal survival and productivity (Kirk, 1994; Pruvost et al., 2019). The experimental results demonstrate a stark contrast in light tolerance between the strains: *C. reinhardtii* CC1690 exhibited an apparent susceptibility to the light intensity tested, aligning with its natural preference for lower light conditions (**Figure 7**). In contrast, *C. pacifica* strains 402 and 403 and *C. sp.* strain 3222 maintained viability even at high light intensities, indicating a robust photoacclimation capacity that could be highly beneficial for cultivation in regions with intense solar exposure.

**Figure 7:**
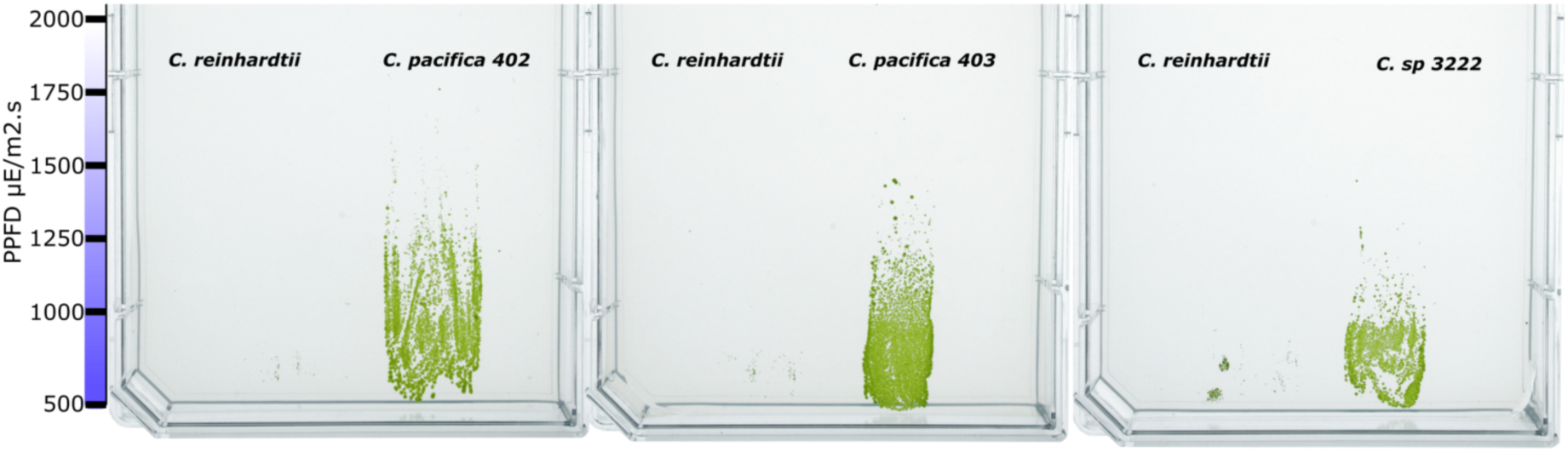
Light Intensity Tolerance of *Chlamydomonas* Strains in HSM Media. The plates demonstrate the growth of *C. reinhardtii* CC1690, *C. pacifica* strains 402 and 403, and *C. sp.* strain 3222 under high light conditions ranging from a minimum of 1000 µE/m²·s to intensities exceeding 2500 µE/m²·s over two days, followed by incubation at 80 µE/m²·s for 5 days. The gradient of light exposure is visually depicted by the diminishing density of algal growth from the bottom (1000 µE/m²·s) to the top of each plate (>2500 µE/m²·s). The comparative analysis highlights the differential tolerance to high light conditions among the strains, with *C. reinhardtii* CC1690 showing notable sensitivity to higher light intensities, while *C. pacifica* strains and *C. sp.* strain 3222 exhibit more robust growth, suggesting a greater resilience to light-induced stress, which may confer an advantage in high-light environments.

This resilience to high light conditions is particularly advantageous in semi-arid regions with intense and consistent solar irradiance. Algae with high light tolerance can utilize abundant sunlight for photosynthesis without the detrimental effects of photoinhibition, thereby maintaining high growth rates and productivity (Cazzaniga et al., 2022; Teoh et al., 2013).

### Cellular and Behavioral Characterization

#### Cell Morphology

Microscopy pictures of *C. pacifica* strains 402 and 403 were captured through differential interference contrast (DIC) microscopy and fluorescence microscopy techniques (**Figure 8**). The DIC images revealed the structural details of the cells, including the presence of two flagella per cell, each with an approximate length of 10 µm. This morphological feature is essential for the cells’ navigation and positioning in their environment, which is crucial for light capture, avoiding unfavorable conditions, and mating.

**Figure 8:**
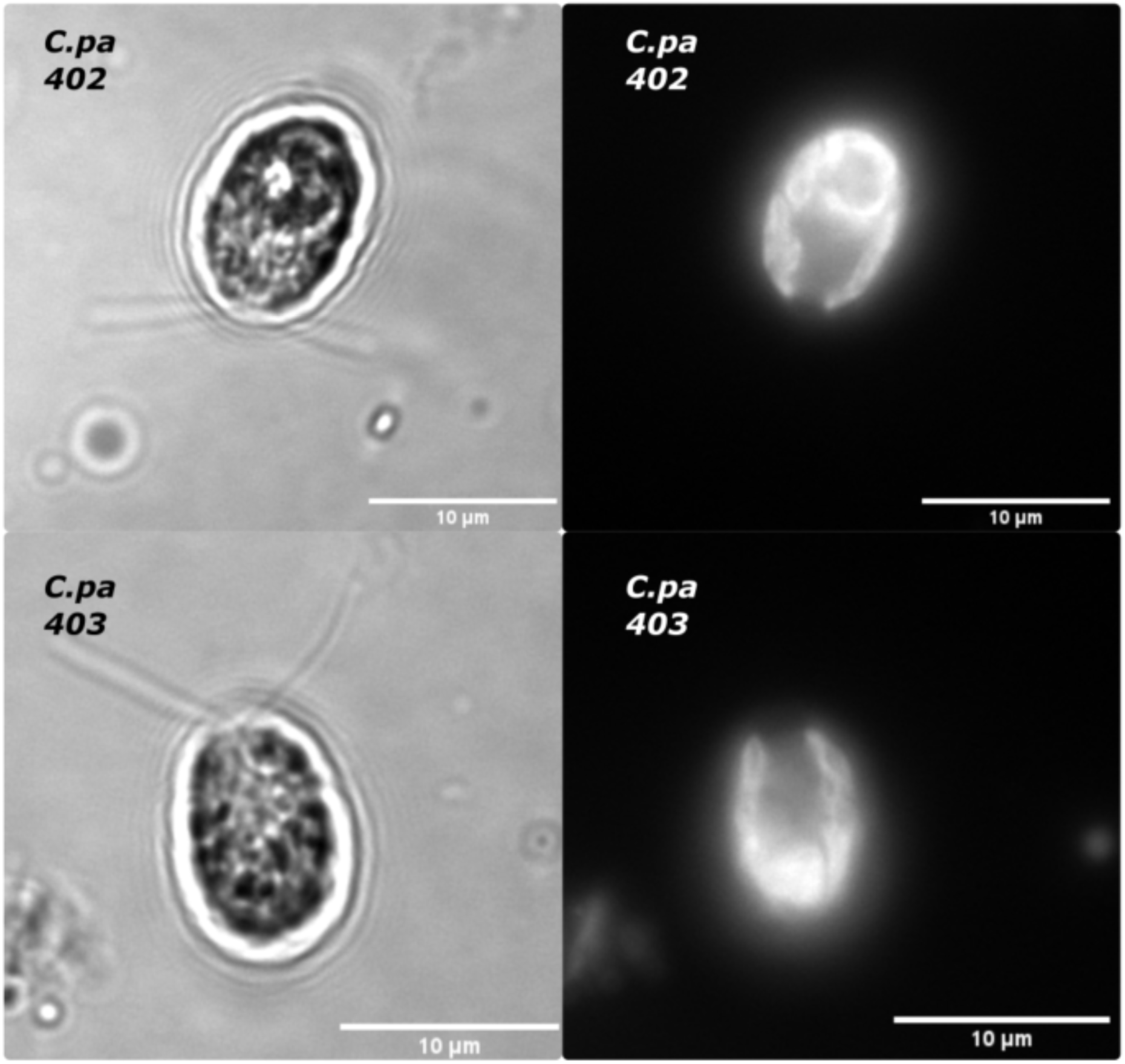
Microscopic analysis of *Chlamydomonas pacifica* strains 402 and 403. The left images display cells of *C. pacifica* strains 402 (top) and 403 (bottom) visualized using Differential Interference Contrast (DIC) microscopy, which reveals detailed cellular structures. The right images correspond to the same cells, highlighting the chlorophyll autofluorescence, indicative of the chloroplasts within the cells. The scale bars represent a length of 10 µm.

The fluorescence microscopy images highlighted the chloroplasts (**Figure 8**). A darker region within the chloroplasts was also observed, likely indicative of a pyrenoid—an organelle linked to carbon fixation (Mackinder et al., 2016). Exploiting *C. pacifica* annotate genome we identified a gene in *C. pacifica* (anno1.g15702.t1) that shows the closest similarity to the *C. reinhardtii* Essential Pyrenoid Component 1 (EPYC1, also known as LCI5) gene, with 52.49% identity (99% coverage and an e-value of 1e-29). EPYC1 is known to associate with Rubisco holoenzymes to form the pyrenoid matrix (Mackinder et al., 2016). These findings suggest that pyrenoids, a common feature in *C. reinhardtii*, may be present in strains 402 and 403 of *C. pacifica*, potentially sharing a similar carbon fixation mechanism as this well-studied species.

#### Motility

The observed motility patterns of *C. pacifica* 402, as depicted in the density graph (**Figure 9; Supplementary Video 1-6**), indicate the species’ adaptive responses to environmental stimuli, highlighting their ecological robustness. The observed range of cell speeds recorded is similar to *C. reinhardtii* (Molino et al., 2022), with cells reaching 150 μm/s. The ability to move strategically allows these algae to seek desirable niches for growth and survival, especially in extreme habitats where resources are scarce or conditions are rapidly changing. This characteristic is particularly advantageous for extremophiles that inhabit such environments because motility facilitates the exploitation of microhabitats that provide refuge from extreme stressors or access to nutrients.

**Figure 9:**
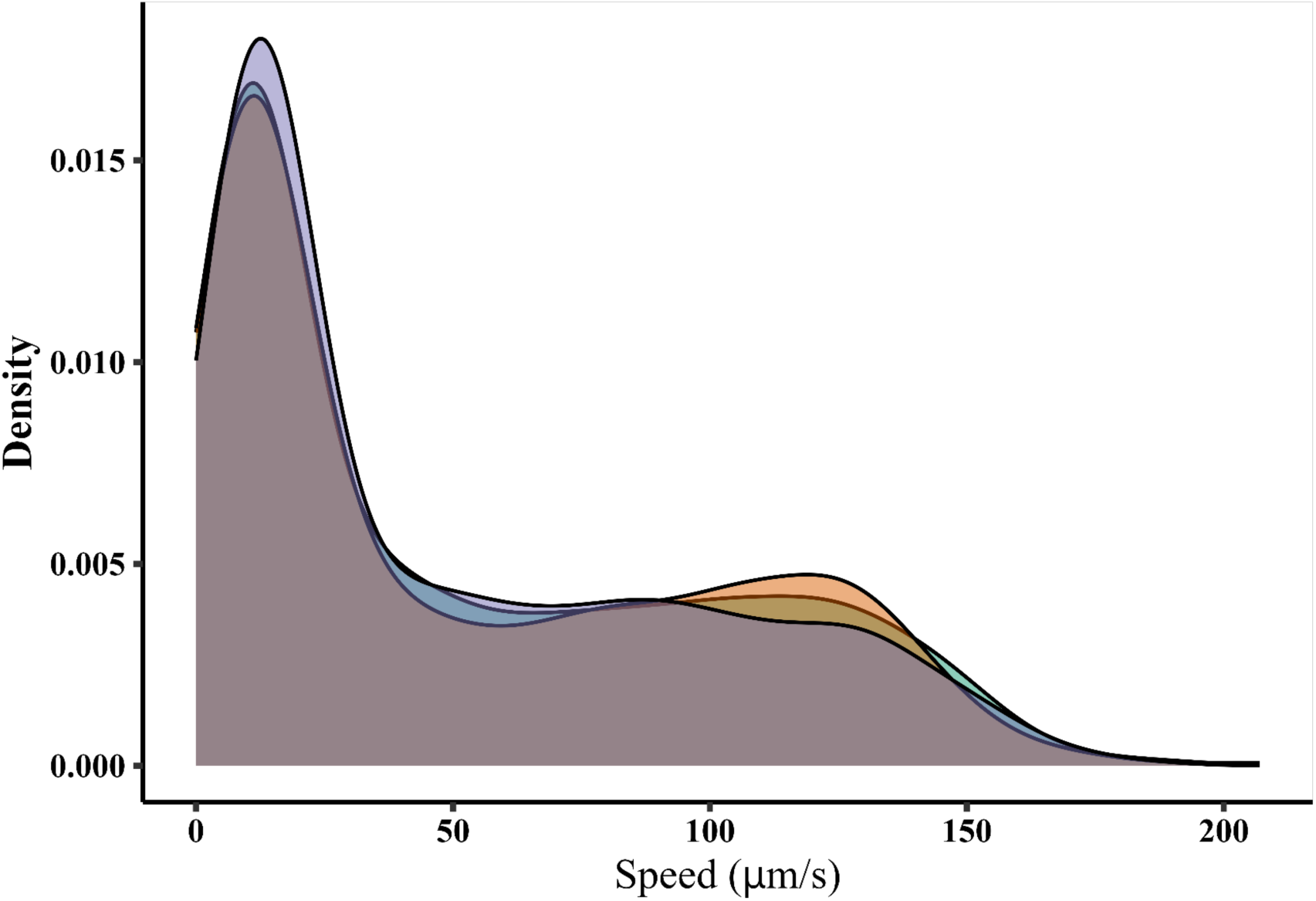
Density Distribution of *C. pacifica* 402 Cell Speed in HSM Media without nitrogen. This graph represents the speed distribution of *C. pacifica* 402 cells, observed under light microscopy. The density curve suggests a range of cellular speeds, with peaks indicating preferred speed ranges, with the fastest cells swimming at 150 μm/s. The plot represents data from three independent biological replicates depicted by each curve.

From a biotechnological standpoint, the motility of algae like *C. pacifica* can be pivotal in industrial applications. In photobioreactors, for instance, active movement can improve the distribution and exposure of algal cells to light, enhancing photosynthetic efficiency and, consequently, biomass productivity (Martinez Carvajal et al., 2024). In applications such as bioremediation, the mobility of algal cells can be crucial for navigating towards environmental pollutants and crucially aiding in biofilm formation (Singh et al., 2006). The motility algae could also be exploited in micromotor drug delivery systems, where extremophile strains could serve as microrobotic platforms to transport drugs to the highly acidic gastrointestinal tract (F. Zhang et al., 2022).

#### Phototaxis

To observe the cell’s phototaxis capacity, we placed *C. pacifica* in HSM without nitrogen and exposed to light from one side of the well. The starving conditions are a stimulus to flagella activity, and the cell, when exposed to light, can use the light cue to display phototaxis (**Figure 10**). We captured two sets of images labeled “White” indicating the white backlight used, and “Chloro” indicating that we used chlorophyll fluorescence to observe cells’ position in the photograph. The cells are evenly distributed across the wells at the initial time (t=0 min). But, after 10 minutes (t=10 min), the cells’ migration away from the light source is evident, congregating at the right corner of the wells, particularly noticeable in the “Chloro” condition.

**Figure 10:**
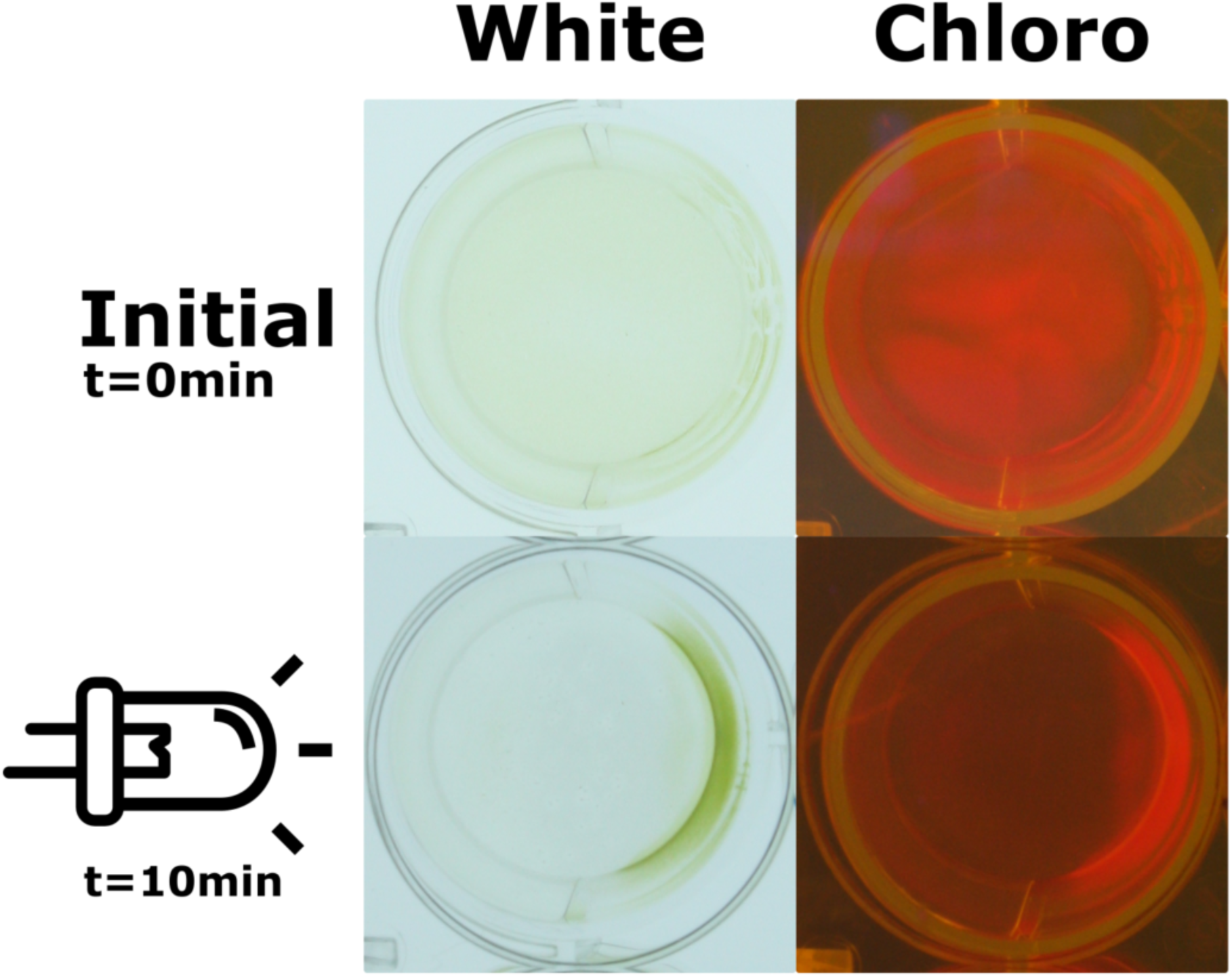
Phototaxis experiment demonstrating *C. pacifica* motility and response to light. The top panel shows strain 402 at the initial moment without light stimuli. The bottom panel shows the phototaxis behavior of the cell after 10 min of exposure to light on the left side of the well. Cells were kept in HSM-N media with a light intensity of 300 uE/m^2^s. Photos were obtained with a white transilluminator (left panel) and a blue transilluminator with an orange light filter (right panel) to observe Chlorophyll pigment fluorescence.

Lower light was present in the right corner. This movement indicates a negative phototactic response of the algae cells, moving away from the light to a more optimal intensity to perform photosynthesis or to avoid detrimental effects from the high amount of light. Phototaxis is a trait common in the species from the core-Reinhardtinia clade, as observed in Volvox, Gonium and *Chlamydomonas reinhardtii* (Leptos et al., 2023). Interestingly, phototaxis can be harnessed to guide cells as microrobots, allowing precise control of their movement. This capability has been demonstrated in *C. reinhardtii*, where phototactic behavior was utilized to direct algae cells loaded with antibiotics for targeted drug delivery onto bacterial cells (Shchelik et al., 2021).

#### Mating

We observed mating behavior between strains *C. pacifica* 402 and 403 (**Figure 11; Supplementary Video 7 and 8**). The observed behavior of *C. pacifica* strains 402 and 403 provides compelling evidence of sexual reproduction. This process plays a crucial ecological role by contributing to genetic diversity and population adaptability (Harris et al., 2009a), but it is also an essential tool for breeding specific traits in a single strain (Fields et al., 2019). *C. reinhardtii* gametes display flagella, which is crucial for mating and is present in *C. pacifica* (**Figure 11A**). As seen in **Figure 11B**, the subsequent release of the cell wall was also detected and is an essential prelude to gamete fusion (Goodenough & Heitman, 2014).

**Figure 11:**
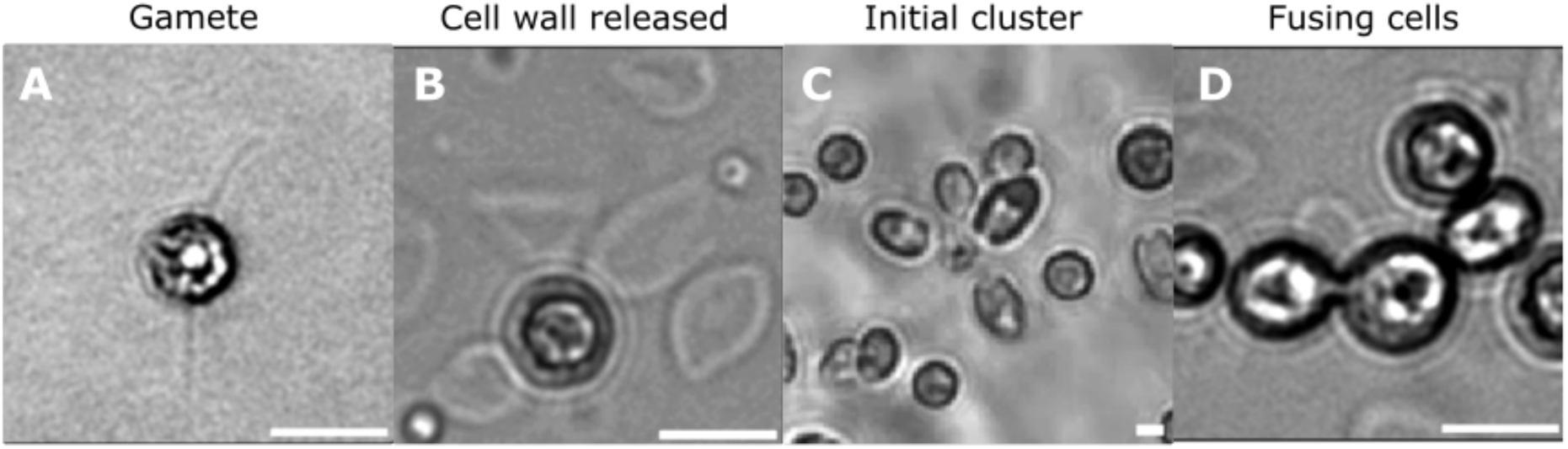
Sequential Stages of Mating in *C. pacifica* Strains 402 and 403. The series of images captures the progressive phases of the mating process. The first image depicts an isolated gamete with flagella ready for mating. The second image shows cells after the release of their cell walls, a preparatory step for gamete fusion. The third image illustrates the initial clustering of cells as they aggregate to form mating pairs. The final image captures a pair of cells in the act of fusion, with the fusion channel visibly formed between them, facilitating the exchange of genetic material. This sequence of events underscores the complex mating behavior exhibited by these algae and provides a visual representation of sexual reproduction in *Chlamydomonas* species. The scale bar indicates 10 um.

As the process unfolds, initial cell aggregation occurs, a behavior fundamental to the formation of mating pairs, as shown in **Figure 11C** and **Supplementary Video 7**. This clustering is not merely a physical congregation but a highly regulated event mediated by pheromones and cellular recognition mechanisms in *C. reinhardtii* (Ferris et al., 2002b). The conclusion of this sequence is the fusion of the mating pair, captured in **Figure 11D** and **Supplementary Video 8**, where the fusion channel is evident, establishing a direct conduit for genetic exchange (Harris et al., 2009a). This event is a pivotal moment in the sexual life cycle of these organisms and is critical for recombination and the introduction of genetic variability (Colegrave, 2002). The ability of *C. pacifica* strains to engage in sexual reproduction has profound biotechnological implications. This behavior facilitates breeding programs, allowing for the selection and crossing of strains to combine desirable traits, and would allow improved biofuel production efficiencies or enhanced stress resilience (Fields et al., 2019). The genetic tractability implied by sexual reproduction makes *C. pacifica* an attractive candidate for such programs, as it enables the manipulation of genetic material to achieve targeted improvements in algal strains, which could significantly advance microalgal biotechnology applications. To date, we have isolated two strains of this newly identified species, representing a relatively low diversity pool. However, the assembled genome provides a valuable foundation for bioprospecting campaigns to identify additional *C. pacifica* strains to expand the genetic diversity pool. Attempts to cross *C. sp.* 3222 with both *C. pacifica* strains did not result in any observed mating behavior, suggesting reproductive isolation between these closely related species.

### Genetic engineering

The capacity for genetic manipulation is a cornerstone in utilizing microorganisms for biotechnological applications. The ability to modify and control the genetic makeup of organisms such as algae enables the development of tailored solutions to address a wide array of challenges, from sustainable production of biofuels to environmental remediation. In this context, *C. pacifica* emerges as a promising candidate due to its amenability to genetic engineering.

We demonstrate the genetic manipulation of *C. pacifica* (**Figure 12**). *C. pacifica* has been successfully modified to express the fluorescent protein mCherry cytosolic and display resistance to zeocin, by nuclear transformation. Also, we successfully showed the genetic modification of *C. pacifica* strain 403 to secrete PHL7 (Sonnendecker et al., 2022), a plastic-degrading enzyme, underscoring this strain’s remarkable potential in the biotechnology field. The clear halos observed around the colonies on hygromycin selective media highlight the strain’s proficiency in protein secretion, a complex trait with diverse applications ranging from industrial processing to environmental cleanup (**see Material and Methods**).

**Figure 12.**
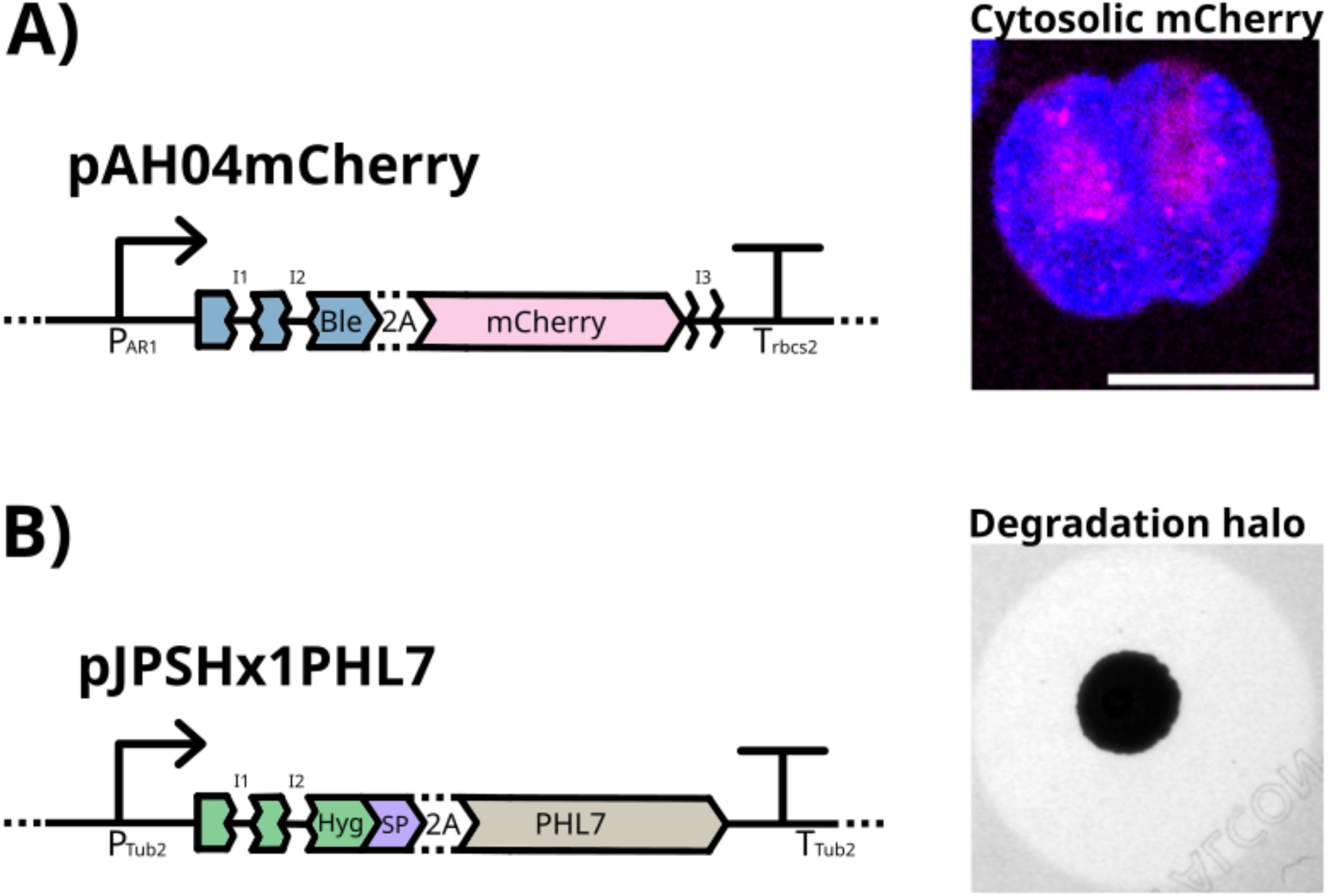
Validation of *Chlamydomonas pacifica* recombinant strains expressing cytosolic mCherry and secreted PHL7 in the opposing mating types. (A) The vector map of the pAH04mCherry plasmid illustrates the construct enabling cytosolic expression of mCherry in *C. pacifica 402* (CC-5697), driven by the *P_AR1_* promoter and *T_rbcS2_* terminator, and including a Bleomycin resistance gene (*Ble*) and a 2A self-cleaving peptide sequence. The confocal microscopy image confirms cytosolic localization of mCherry fluorescence (magenta) in the cytosol alongside chlorophyll autofluorescence (blue) in the chloroplast, with a scale bar representing 10 µm. **(B)** The vector map of the pJPSHx1PHL7 plasmid shows the construct for the secretion of PHL7 in *C. pacifica 403* (CC-5699), under the control of the *P_Tub2_* promoter and *T_Tub2_* terminator, including a Hygromycin resistance gene (*Hyg*), a signal peptide (SP), and a 2A peptide. The plate assay image demonstrates a degradation halo on HSM acetate agar containing zeocin (15 µg/mL) and supplemented with 0.5% Impranil® DLN, indicating enzymatic degradation of the plastic dispersion. These results confirm that *C. pacifica* can be genetically engineered in both mating types to express a cytosolic fluorescent protein (*C. pacifica 402*) or secrete an active plastic-degrading enzyme (*C. pacifica 403*).

In this study, we present further evidence of the successful mating between *Chlamydomonas pacifica* strains 402 and 403, reinforcing the sexual reproduction capabilities within these algal strains. The image captured post-mating shows progeny colonies growing on a selective medium containing hygromycin (30 µg/mL) and zeocin (15 µg/mL), supplemented with 0.5% Impranil (**Figure 13**). The dual resistance to both antibiotics in the progeny is an indication of genetic material exchange between the two parent strains, as each parent harbored a distinct resistance marker: strain 402 contained the pJPSHx1PHL7 vector for hygromycin resistance, and strain 403 carried the pAH04mCherry vector for zeocin resistance. The survival and growth of these progeny colonies under stringent selection conditions corroborate the mating event and confirm the inheritance and functional expression of both antibiotic-resistance genes. This demonstrates the successful combination of genetic traits through sexual reproduction.

**Figure 13:**
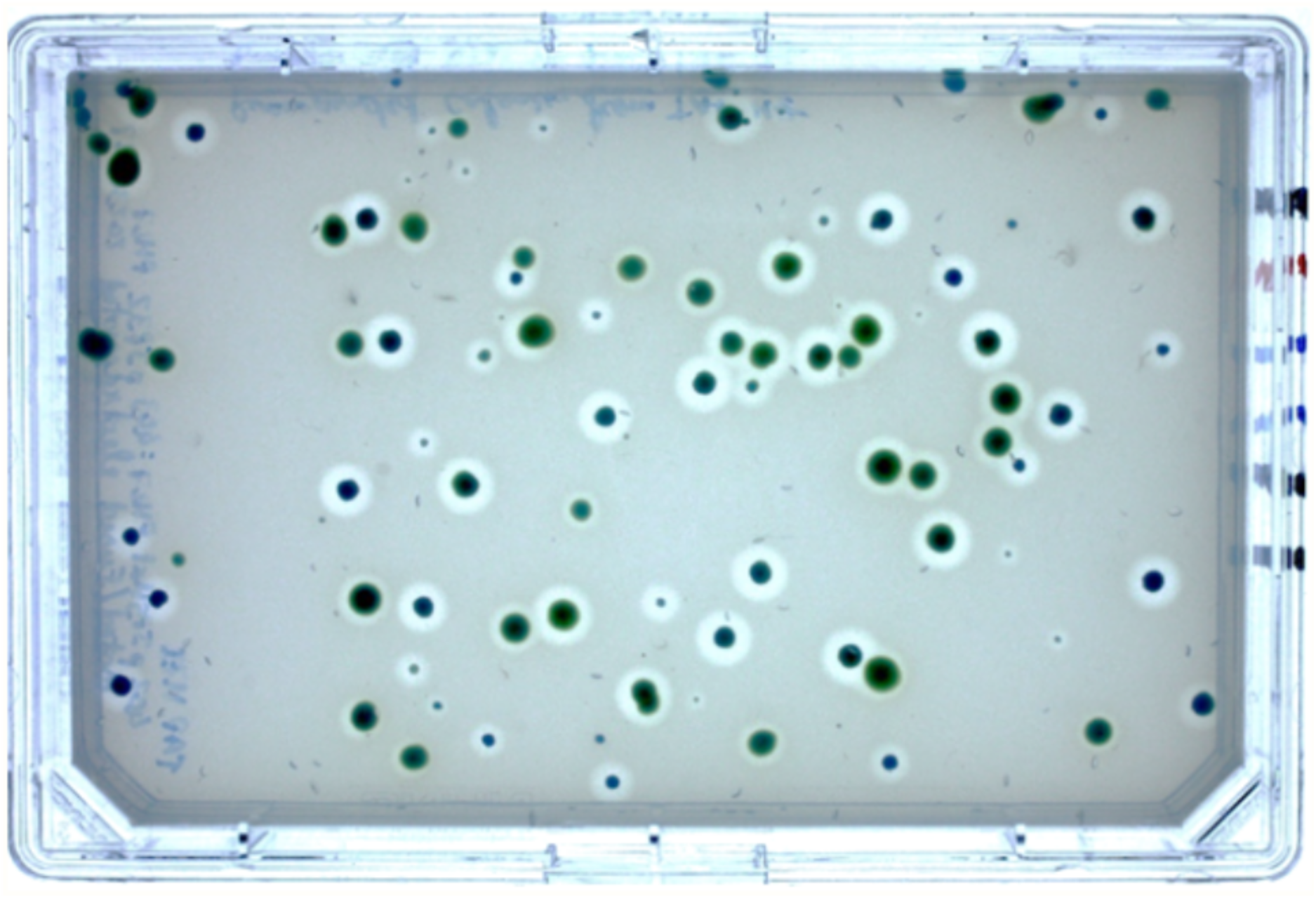
Selection of Progeny from Mating of *Chlamydomonas pacifica* Strains 402 (Hygromycin Resistant) and 403 (Zeocin Resistant). This image showcases the progeny resulting from the mating of *C. pacifica* 402, which harbors the pJPSHx1PHL7 vector conferring hygromycin resistance and the ability to secrete PHL7, with *C. pacifica* 403, carrier of the pAH04mCherry vector that confers zeocin resistance and encodes for cytosolic expression of mCherry. The selection plate contains hygromycin (30 µg/mL) and zeocin (15 µg/mL), along with 0.5% Impranil, ensuring that only progeny inheriting both resistance genes and the associated PHL7 secretion capability survive. The presence of green colonies amidst the selective agents indicates successful mating and genetic exchange between the two strains, leading to a new generation of algae with combined traits for antibiotic resistance and enzymatic activity.

### Lipid profile

We performed a comprehensive lipidomic profile of *C. pacifica*, derived from both negative and positive ionization modes of mass spectrometry (**Table 1 and 2**). These tables encapsulate the diverse lipid species detected within the algal strain, with a rich array of triacylglycerols (TAGs) pertinent to biofuel applications. Additionally, we observed some relevant nutraceuticals and dietary supplements, given the health-promoting potential of the fatty acids identified in these lipid classes. We further discuss each possible application below.

**Table 1:**
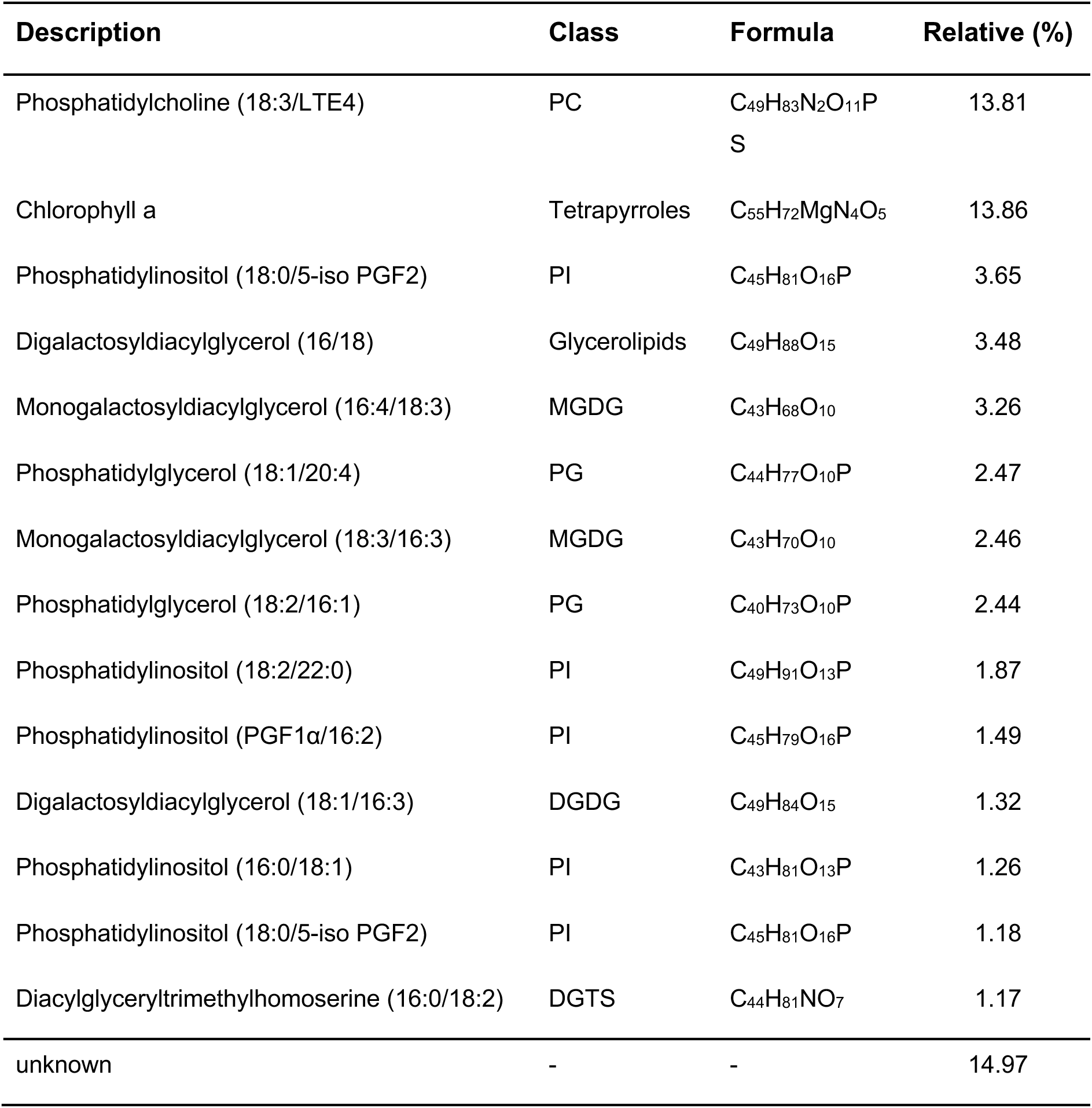
Lipidomic profile of *Chlamydomonas pacifica*: identification, classification, and relative abundance of lipid species in negative mode analysis.

#### Triacylglycerol Composition and Biofuel Potential

Lipidomic analysis of *C. pacifica*, utilizing both positive and negative ionization modes, has revealed a lipid profile rich in triacylglycerols (TAGs), which are particularly interesting for biofuel applications. Specifically, the positive ionization mode data elucidated a series of TAGs, comprising glycerol bound to three fatty acids, making them highly reduced forms of carbon and, thus, energy-dense molecules suitable for biofuel production (**Table 1 and 2**).

Among the detected lipids, several TAGs were present with chain lengths of 16 and 18 carbons, including unsaturated versions. These fatty acids can be converted to biodiesel, and *C. pacifica* can be improved to maximize lipid yield, by genetic engineering or breeding programs. The extraction process can also be enhanced to improve the economic feasibility of biofuel production. Additionally, metabolic engineering strategies could increase the proportion of desirable fatty acids, such as increasing the saturation level to improve biodiesel stability or tweaking the carbon chain length to favor biodiesel properties (Ibáñez-Salazar et al., 2014; Lu et al., 2008).

#### Fatty Acid Composition and Health-Promoting Potential

The comprehensive lipidomic profile of *C. pacifica* has uncovered a spectrum of lipid species with significant potential for nutraceuticals and dietary supplements. Notably, the analysis identified an array of TAGs and phospholipids with fatty acid compositions that benefit human health (**Table 1 and 2**). Among the lipid compounds identified, the presence of TAGs containing omega-3 and omega-6 fatty acids, such as linoleic acid (18:2) and α-linolenic acid (18:3) (**Table 2**). These polyunsaturated fatty acids (PUFAs) are essential components of the human diet as well, as they are precursors to eicosapentaenoic acid (EPA) and docosahexaenoic acid (DHA), which play critical roles in maintaining cardiovascular health and cognitive function (Swanson et al., 2012). The analysis also revealed phospholipids like phosphatidylcholines (PCs) and phosphatidylinositols (PIs) (**Table 1**), which are integral to cell membrane structure and have been associated with health benefits, including liver health and cognitive improvements (Küllenberg et al., 2012).

**Table 2:** Lipidomic profile of *Chlamydomonas pacifica*: identification, classification, and relative abundance of lipid species in positive mode analysis.

| Description | Class | Formula | Relative (%) |
| --- | --- | --- | --- |
| Chlorophyll a | Tetrapyrroles | C<sub>55</sub>H<sub>72</sub>MgN<sub>4</sub>O<sub>5</sub> | 6.59 |
| Triacylglycerol (16:0/18:2/18:3) | TG | C<sub>55</sub>H<sub>96</sub>O<sub>6</sub> | 2.64 |
| Triacylglycerol (16:0/18:1/18:3)[iso6] | TG | C<sub>55</sub>H<sub>98</sub>O<sub>6</sub> | 2.48 |
| Galactosylceramide(d18:2/20:1) | GalCer | C<sub>44</sub>H<sub>81</sub>NO<sub>8</sub> | 2.24 |
| Triacylglycerol (16:0/18:0/18:2)[iso6] | TG | C<sub>55</sub>H<sub>102</sub>O<sub>6</sub> | 2.07 |
| Diacylglyceryltrimethylhomoserine (16:0/18:2) | DGTS | C<sub>44</sub>H<sub>81</sub>NO<sub>7</sub> | 1.97 |
| Triacylglycerol (16:0/18:1/18:2)[iso6] | TG | C<sub>55</sub>H<sub>100</sub>O<sub>6</sub> | 1.89 |
| Phosphatidic acid (22:5/18:3) | PA | C<sub>43</sub>H<sub>69</sub>O<sub>8</sub>P | 1.69 |
| Triacylglycerol (16:0/17:2/17:2)[iso3] | TG | C<sub>53</sub>H<sub>94</sub>O<sub>6</sub> | 1.56 |
| Diacylglycerol(16:0/18:2) | DG | C<sub>49</sub>H<sub>88</sub>O<sub>15</sub> | 1.47 |
| Monogalactosyldiacylglycerol (18:3/16:3) | MGDG | C<sub>43</sub>H<sub>70</sub>O<sub>10</sub> | 1.42 |
| Triacylglycerol (16:0/16:1/18:2)[iso6] | TG | C<sub>53</sub>H<sub>96</sub>O<sub>6</sub> | 1.35 |
| GalCer(d18:1/20:0) | GalCer | C<sub>44</sub>H<sub>85</sub>NO<sub>8</sub> | 1.25 |
| phosphatidylcholine (ladderane-C6/ladderane-C8) | PC | C<sub>46</sub>H<sub>80</sub>NO<sub>6</sub>P | 1.24 |
| unknown | - | - | 9.37 |

Identifying these lipid classes in *C. pacifica* offers exciting possibilities for developing algae-based nutraceuticals and dietary supplements. The cultivation of microalgae as a source of PUFAs and other bioactive lipids is a growing area of interest due to the sustainability of algae as a resource and the potential health benefits they offer (Diaz et al., 2023).

## Conclusion

Microalgae hold tremendous potential for the bioeconomy and can be used to support several human endeavors. The newly isolated species *C. pacifica* has demonstrated interesting traits that make it a chassis for bioprocessing using algae. It harbors the common characteristics desirable in an industrial strain, such as the capacity to grow in a simple medium, and high tolerance to abiotic stressors common in non-arable arid regions such as high temperature, salinity, and light. However, there are also more niche traits, such as the high pH tolerance bound to support farming this algae. The mating capacity between strains 402 and 403 and the proven potential for genetic manipulation exemplify the organism’s suitability for advanced biotechnological processes and strategies to manipulate and improve the strain to different goals. These traits and the alga’s genetic tractability present *C. pacifica* as an excellent candidate for future metabolic engineering endeavors aimed at sustainable biotechnological solutions.

## Material and Methods

### Isolation and Characterization

Water samples were collected from a pond in the Biology field station of UCSD (32° 53 ‘09”N, 117° 13’ 48”W) and plated on High-Salt Media (HSM) media (**Supplementary Table 3a**). Colonies were repeatedly streaked on the same medium to obtain pure colonies as confirmed by a single morphology and size when examined directly using a light microscope. Throughout the year 2023, this region exhibits an average monthly air temperature fluctuating between 10°C and 25°C, alongside solar radiation averages that range from 128 W/m^2^ to 291 W/m^2^ monthly, according to the California Irrigation Management Information System (CIMIS). The opposing mating type strains were deposited at the Chlamydomonas Collection under strain numbers CC-5697 (402) & CC-5699 (403). Throughout the text, we maintained the names *C. pacifica* 402 for CC-5697 and *C. pacifica* 403 for CC-5699.

### Growth Conditions

The cultures were grown in various media, all described in **Supplementary Table 3a-d**, at 25-28 °C under constant illumination of 80 μmol photons/m^2^s and agitation of 150 rpm in 50mL cultures unless stated otherwise. To obtain the growth curve, 160 uL media inoculated with samples added in 96 well plates and read in the Infinite® M200 PRO plate reader (Tecan, Männedorf, Switzerland) from 6 biological replicates for each strain and condition studied. The cultures were grown at 30 °C under constant illumination of 80 μmol photons/m^2^s at 800 rpm on a rotary microplate shaker.

### Genomic DNA and RNA Extraction

Cells were grown in HSM media and spun down, flashed frozen in liquid nitrogen and sent frozen to UCDavis genome center. The DNA was sequenced in the Nanopore sequencer PromethION for long reads and in Illumina NovaSeq 6000 for short reads. The RNA samples were extracted from cells grown until stationary phase photosynthetically in HSM media, mixotrophically in Tris-Acetate-Phosphate (TAP) media, heterotrophically in TAP media, and photosynthetically in high salt on the D2-15 media supplemented with 15g/L of Na_2_CO_3_. The RNA was extracted by pelleting cells by centrifugation at 3,000 × g for 1 minute at 4°C. The pellet was then resuspended in 0.25 mL lysis buffer containing 50 mM Tris-HCl (pH 8.0), 200 mM NaCl, 20 mM EDTA, 2% SDS, and Proteinase K (0.5 mg/L). The lysate was incubated at 70°C for 2 minutes to ensure thorough lysis and was then mixed with 2 mL of TRIzol reagent. The mixture was incubated at room temperature for 5 minutes, followed by adding 0.5 mL of chloroform, which was vigorously shaken for 15 seconds. After a 5-minute incubation at room temperature, the sample was centrifuged at 12,000 × g for 15 minutes at 4°C. The aqueous phase was carefully transferred to a new tube, and RNA was precipitated by adding 1 mL of isopropanol, followed by a 10-minute incubation at 4°C. The RNA was pelleted by centrifugation at 12,000 × g for 10 minutes at 4°C, and the supernatant was discarded. The RNA pellet was washed by resuspension in 2 mL of 75% ethanol, briefly vortexed, and centrifuged at 7,500 × g for 5 minutes at 4°C. The supernatant was discarded, and the RNA pellet was air dry for 5–10 minutes. The RNA pellet was resuspended in 25 µL of RNase-free water, 3.5 µL of 10X DNase buffer, and 5 µL of DNase were added, and the sample was incubated at room temperature for 1–2 hours. Finally, 3 µL of EDTA was added to the RNA solution, and the mixture was incubated at 65°C for 10 minutes to inactivate the DNase. The RNA samples were submitted to the UCSD IGM Genomics Center.

### Genome Sequencing and Annotation

Long DNA reads were produced using Oxford Nanopore sequencing on the PromethION platform. These long reads were cleaned, trimmed, and assembled with Flye version 2.9.3, using default program parameters except for an estimated genome size of 120m (Kolmogorov et al., 2019). This assembly produced 14 contigs longer than one million nucleotides, with an N50 of 484,211. Illumina sequencing was performed using the Illumina NovaSeq 6000 platform to improve local misassemblies in the genome. These short reads were cleaned and trimmed using fastp version 0.23.1 with default parameters (Chen et al., 2018). Illumina reads were given as input to Pilon version 1.24, and all downstream analyses were performed on this polished genome (Walker et al., 2014).

Additional RNA sequencing was performed from four samples using the Illumina NovaSeq 6000 platform to support gene model predictions. These transcriptomic reads were cleaned using fastp and then mapped to the genome using STAR version 2.7.4a (Dobin et al., 2013). Mapped transcriptomic reads, the AUGUSTUS model for *Chlamydomonas reinhardtii*, and the OrthoDB version 10 set of Viridiplantae proteins were given to Braker v. 3.0.6 to predict gene models (Gabriel et al., 2024; Hoff & Stanke, 2019; Kuznetsov et al., 2023). These gene models were annotated using the software eggNOG-mapper version 2 eggNOG version 5.0 database (Cantalapiedra et al., 2021; Huerta-Cepas et al., 2019). Genomic assembly and gene model predictions were evaluated for completeness using BUSCO version 5.5.0 in both genome and protein modes, compared to a set of conserved chlorophyte genes provided in OrthoDB (Kuznetsov et al., 2023; Seppey et al., 2019).

### Phylogenetic Analysis

For constructing the phylogenetic tree, we considered all species under the Chlorophyta taxon whose gene annotations were made available on the NCBI genome platform (https://www.ncbi.nlm.nih.gov/datasets/genome). We incorporated 1,519 genes from the Chlorophyta odb10 dataset, as identified by the Benchmarking Universal Single-Copy Orthologs (BUSCO) initiative (Manni et al., 2021). For each BUCSO gene, we identified its orthologue in each species by employing the protein mode of BLAST and selected the gene with the lowest e-value (Johnson et al., 2008). Subsequently, we performed multiple-sequence alignment using MUSCLE and generated the corresponding gene trees using CLUSTALW2 (Edgar, 2004; Larkin et al., 2007). Finally, the species tree was estimated from the gene trees using ASTRAL-III (C. Zhang et al., 2018).

### Lipidomics

Pellet samples from algal cultures were prepared through lyophilization and resuspended with ribitol as an internal standard for consistent quantification. Following the adapted protocol from Hollin et al. (2022), a biphasic extraction using a methyl tert-butyl ether and methanol solution facilitated the separation of metabolites, which were then analyzed for lipidomics at the UC Riverside Metabolomics Core Facility using a G2-XS quadrupole time-of-flight mass spectrometer (Hollin et al., 2022). Data analysis incorporated advanced software like Progenesis Qi for peak identification, normalization, and annotation, employing various databases and proprietary tools for comprehensive metabolite profiling. The detailed methodological procedure is available at Supplementary Material (see **Supplementary Method** section).

### Strain profile

#### Carbon and Nitrogen Assimilation

To evaluate carbon and nitrogen assimilation, HSM and HSM without nitrogen agar plates were prepared using Thermo Fisher Scientific’s Thermo Scientific™ Nunc™ OmniTray™ Single-Well Plate (Catalog number: 242811). The composition of the media is outlined in the Supplementary Material (**Supplementary Table 3a-d**). Each plate containing approximately 35 ml of 1.5% agar medium was prepared. Subsequently, 1 ml of algae culture was evenly spread across the surface of the plate. Once the cells had dried to the corners of the plate, 500 ul of a 1% agarose solution containing HSM with varying nitrogen sources or different carbon source solution was gently dispensed onto the surface, creating a small mound above the agar medium. The carbon source plates were left on the lab bench in low light of approximately 8 µmol photons m^2^ s^-1^, and nitrogen source plates were grown photosynthetically in 80 µmol photons/m^2^s^1^ of light.

#### pH Tolerance

To compare the strain’s capacity to grow in different pH and salinity, Nunc® OmniTray (Merck KGaA, Darmstadt, Germany) was prepared with agar HSM. The pH gradient was created by adding and spreading 60 µL of 3.3M H_3_PO_4_ on one side of the plate, followed by adding 60 µL of 10M KOH on the opposite side. The plates were left to equilibrate for 2 days, so the pH gradient was formed, and samples were spread across the pH gradient in parallel sections of the plate for pH tolerance capacity (**Supplementary** Figure 2).

#### Salt Tolerance

To compare the strain’s capacity to grow in different salinity, Nunc® OmniTray (Merck KGaA, Darmstadt, Germany) were prepared with agar HSM media with 150 mM NaCl. The salt gradient was created by removing a section of 1.5 cm in one of the sides of the plate and adding a melted 1.5% agarose solution of 5M NaCl. To further compare the strain’s capacity to grow in different salinities, growth curves were prepared in HSM with varying additional concentrations of NaCl, grown in 96 well plates, shaken at 800 rpm with continuous light at 80 µmol photons m^2^ s^-1^ from white neutral LEDs. Growth was monitored by optical density measured at 750 nm in the Infinite® M200 PRO plate reader (Tecan, Männedorf, Switzerland).

#### Temperature Tolerance

Cells were grown in HSM acetate until the stationary phase to compare the strains’ capacity to temperature and exposed to different temperature values, ranging from 37.5°C to 44.5°C. The cells were exposed for 24 hours and plated on an agar HSM plate to recover surviving cells. Briefly, 150 µL of culture was added to 96 well PCR plates, and 5 µL samples were taken to an HSM acetate plate and placed to grow to be used as an initial culture condition. The rest of the samples were sealed with a PCR plate seal. The plates were incubated in a 96-well T100 thermal cycler (Biorad, Hercules, CA, USA) using fixed temperature gradients for each row. After 24 hours of incubation, the cells were resuspended by pipetting, and a 5 µL sample was added to an HSM acetate plate. After the culture spots were dried, the plates were placed in growing conditions. After 5-7 days, images were taken from each plate.

#### Light Tolerance

To assess the light tolerance of different strains, cultures were grown to stationary phase in either HSM acetate or TAP media. Subsequently, 200 µL of each culture was uniformly spread along the length of an OmniTray™ Single-Well Plate (Catalog number: 242811) containing HSM agar. *Chlamydomonas reinhardtii*, known for its high-light sensitivity, served as a control and was tested in parallel to the test strains (Erickson et al., 2015; Virtanen et al., 2021). The plates were then positioned 9 cm away from a 1000W Mastiff GrowL® LED grow light, utilizing both blue and red LEDs to simulate high-light conditions. To create a gradient of light exposure, plates were aligned so that portions were directly beneath the LEDs while others extended beyond the light’s direct reach. Protective shading was placed in one of the corners of each plate to ensure survival of some cells as initial inoculum control. The photosynthetic photon flux density (PPFD) was measured across the plate surface at 5 mm intervals and repeated thrice. The resulting data, excluding the initial shadowed area and the intense central light zone of the panel, depicted a linear PPFD gradient, as detailed in **Supplementary** Figure 3.

#### Phototaxis

To investigate phototaxis in *Chlamydomonas pacifica* 402, cells were cultured on HSM agar plates for 5-7 days. Subsequently, the mature cells were harvested and resuspended in nitrogen-free HSM media to trigger gametogenesis and enhance flagellar movement. The prepared cells were then distributed into a 6-well plate and positioned before a light source, allowing them to exhibit phototactic behavior for 10 minutes. Photographic documentation was performed using a Canon EOS camera, capturing images at the start (t=0 min) and after 10 minutes of light exposure from a 15 W fluorescent white light, placed 2 cm from the 6-well plate. To visualize the cells, photographs were taken against both a white backdrop to highlight their green pigmentation and a blue backdrop with an orange filter to detect chlorophyll fluorescence, providing a view of the cells’ phototactic response.

#### Motility

To measure cell movement speeds, we followed the method described in Molino et al., (2022), with a few modifications. Cells were grown in HSM acetate agar plates for 5 days, scraped, and resuspended in HSM media without the nitrogen source. The cells were kept in the dark until testing for approximately 10-15 min. The slides were prepared using glass slides with Frame-Seal™ Slide Chambers, 15 × 15 mm, 65 μL, and sealed with a cover slip. The cells were observed with a Nikon microscope equipped with a Nikon DS-Qi2 camera, and the recorded video was analyzed with ImageJ’s plugin TrackMate (6.01) (Tinevez et al., 2017). The generated motility results were performed in a chamber, allowing cells to swim in all directions, including on the axis parallel to the camera (Z-axis). Measured speeds are only relative to the X and Y axis vectors. A more detailed protocol can be found at dx.doi.org/10.17504/protocols.io.bsw5nfg6. R version 3.6.3 (2020-02-29) running in the RStudio v1.2.5042 IDE was used to import and process data and generate the plots as a density plot.

### Assembly of transformation vectors

All restriction enzymes utilized in this study were sourced from New England Biolabs (Ipswich, MA, USA). These constructs were developed using the pBlueScript II + KS (pBSII) backbone. The vector pAH04mCherry was previously described by Molino et al., (Molino et al., 2018), and it contains genetic elements from *C. reinhardtii*. The vector pJPSHxPHL7 was assembled using genetic elements from the assembled genome from *C. pacifica* 402. We identified the β-Tubulin A 2 gene and its structures. The vector was then assembled using the promoter, introns one and two, and the 3’UTR region of the gene. We added the hygromycin resistance marker codon optimized to *C. reinhardtii* followed by the foot-and-mouth-disease-virus 2A self-cleavage peptide (FMDV-2A), the signal peptide from Sadp1 (Molino et al., 2018) from *C. reinhardtii* and the codon-optimized sequence for PHL7 (Sonnendecker et al., 2022). The map can be observed in **Supplementary** Figure 4, and the vector sequence is available at 10.5281/zenodo.14060701.

### Transformation

Transformation of *C. pacifica* was conducted using electroporation, following the protocol outlined by Molino et al., (Molino et al., 2022). In summary, cells were cultured to reach mid-logarithmic growth phase (3–6×10^6^ cells/mL) in HSM acetate medium, maintained at 25 °C with constant light exposure (80 μmol photons/m^2^s) and shaking at 150 rpm. The cells were then concentrated by centrifugation at 3000 g for 10 minutes and resuspended to a density of 3–6×10^8^ cells/mL using the MAX Efficiency™ Transformation Reagent for Algae (Catalog No. A24229, Thermo Fisher Scientific). For the electroporation, a mixture of 250 μL of this cell suspension and 1000 ng of a vector plasmid digested with XbaI and KpnI enzymes was chilled on ice for 5–10 minutes within a 4-mm width cuvette compatible with the Gene Pulser®/MicroPulser™ system (BioRad, Hercules, CA). The cell-DNA mixture was then electroporated using the GenePulser XCell™ device (BioRad, Hercules, CA), applying a pulse of 2000 V/cm for 20 microseconds. Following electroporation, the cells were transferred into 10 mL of HSM acetate medium and incubated under gentle agitation (50 rpm) in ambient light conditions (around 8 μmol photons/m^2^s) for an 18-hour recovery period. Post-recovery, the cells were again concentrated by centrifugation, resuspended in 600 μL of HSM acetate medium, and evenly spread onto two HSM acetate agar plates, each containing different antibiotics according to the vectors used. The zeocin (Zeo) concentration was 10 μg/mL and hygromycin (Hyg) was 30 μg/mL. The plates were then incubated under a light intensity of 80 μmol photons/m^2^s at 25 °C until visible colonies developed.

The pJPSHx1PHL7-transformed cells were plated onto TAP agar plates containing 0.75% (v/v) Impranil® DLN, a colloidal polyester-PUR dispersion, and 30 μg/mL hygromycin (Liu et al., 2021). These selective media plates were prepared by making 1 L of TAP media with agar and autoclaving to sterilize. Once the media had cooled to 55°C, hygromycin was added to a final concentration of 30 μg/mL, and 7.5 mL of Impranil® DLN was added to achieve 0.75% (v/v). After thoroughly mixing, the media was poured into Petri dishes and allowed to solidify.

### Mating

The mating assays were performed in line with the methodology outlined by (Findinier, 2023). Initially, recombinant *Chlamydomonas pacifica* strains CC-5697 harboring the pJPSHx1PHL7 (Hyg) vector and CC-5699 harboring the pAH04mCherry vector (Zeo) were cultured on HSM acetate agar plates for 7 days. Post-cultivation, the cells were harvested using a sterile loop, resuspended in nitrogen-depleted HSM media, and subjected to light exposure (80 μmol photons/m^2^s) overnight without agitation. Each strain possessed a distinct antibiotic resistance marker, with one strain being resistant to zeocin and the other to hygromycin. Following light exposure, the strains were combined and incubated in darkness for at least 24 hours without disturbance. After this dark incubation phase, the cells were centrifuged down at 2000g for 5 min, resuspended in fresh media, and placed under standard growth conditions to facilitate the emergence of daughter cells. After 2-3 days, these cells were then plated on selective media containing both antibiotics, enabling the identification and selection of progeny cells that inherited resistance markers from both parent strains.

### Data analysis

R version 3.6.3 (2020-02-29) running in the RStudio v1.2.5042 IDE was used to import and process data, generate the statistical summary, and generate the motility plots. The growth curve was performed using Microsoft Excel 365 (Version 16.0, Microsoft Corporation, Redmond, WA, USA).

## Data Availability

Raw sequencing data and whole genomic assembly for this project is available at the National Center for Biotechnology Information through BioProject PRJNA1151728. Assembled contigs, gene models, and annotations for the *C. pacifica* genome are available on figshare (10.6084/m9.figshare.26824903). All supplemental materials are also included in the figshare link. The supplementary videos are available at 10.5281/zenodo.13626573. The vectors map and sequence are available at 10.5281/zenodo.14060701. The raw images, codon usage table and lipidomics file are available at 10.5281/zenodo.14061071.

## Authors Contributions

**JVDM**: Conceptualization, Data curation, Formal analysis, Investigation, Methodology, Visualization, Writing – original draft, Writing – review & editing

**AO**: Data curation, Formal analysis, Software, Visualization, Writing – review & editing

**HS**: Data curation, Formal analysis, Software, Visualization, Writing – review & editing

**KK**: Investigation, Methodology, Visualization, Writing – review & editing

**BS**: Investigation, Methodology, Writing – review & editing

**CJD**: Investigation, Writing – review & editing

**AG**: Data curation, Investigation, Formal analysis, Software, Visualization, Writing – review & editing

**LJJ:** Investigation, Writing – review & editing

**YT-T**: Writing – review & editing

**NH**: Investigation, Writing – review & editing

**AB**: Investigation, Writing – review & editing

**HJ**: Investigation, Writing – review & editing

**FF**: Investigation, Writing – review & editing

**RS**: contributed to drafting and revising the original manuscript and secured funding for the research.

**SM**: contributed to drafting and revising the original manuscript and secured funding for the research.

## Acknowledgment

The authors thank Jennifer Santini and Marcy Erb (UCSD School of Medicine Microscopy Core, supported by NINDS P30NS047101 grant) for help with confocal microscopy. This publication includes data generated at the UC San Diego IGM Genomics Center utilizing an Illumina NovaSeq 6000 purchased with funding from a National Institutes of Health SIG grant (#S10 OD026929). The lipidomics data were generated at the Metabolomics Core facility at the University of California, Riverside. The icon was made by Pixel Perfect, Freepik, and Uniconlabs from www.flaticon.com.

