## Supplementary Material for "Description of a novel extremophile green algae, *Chlamydomonas pacifica*, and its potential as a biotechnology host"

#### Supplementary Figures

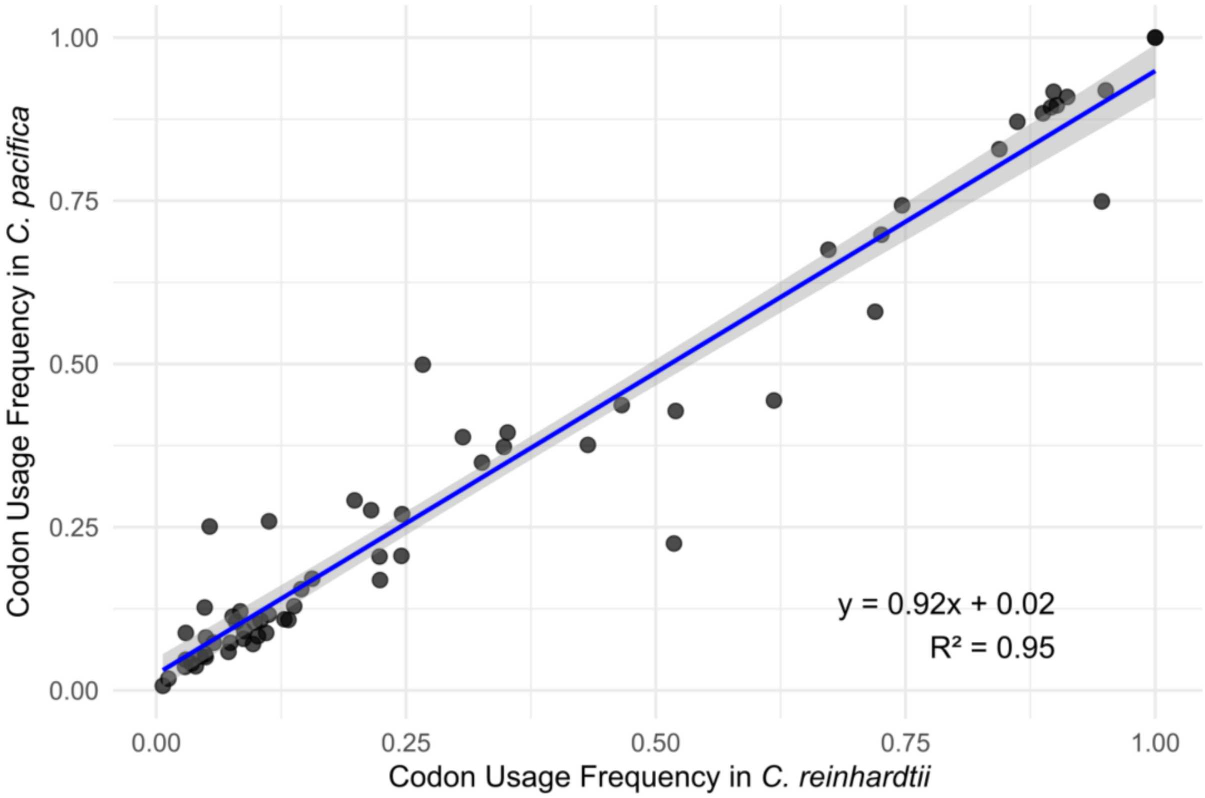


**Supplementary Figure 1**: Comparison of codon usage frequencies between *Chlamydomonas pacifica* and *Chlamydomonas reinhardtii*. The codon usage frequencies were plotted on a scatter plot, with frequencies for *C. reinhardtii* on the x-axis and *C. pacifica* on the y-axis. The plot demonstrates a high degree of similarity in codon usage bias, as shown by the near-linear relationship between the two species' codon usage frequencies. The trend suggests conserved codon preferences between the species, indicative of potential shared translational machinery or evolutionary constraints.

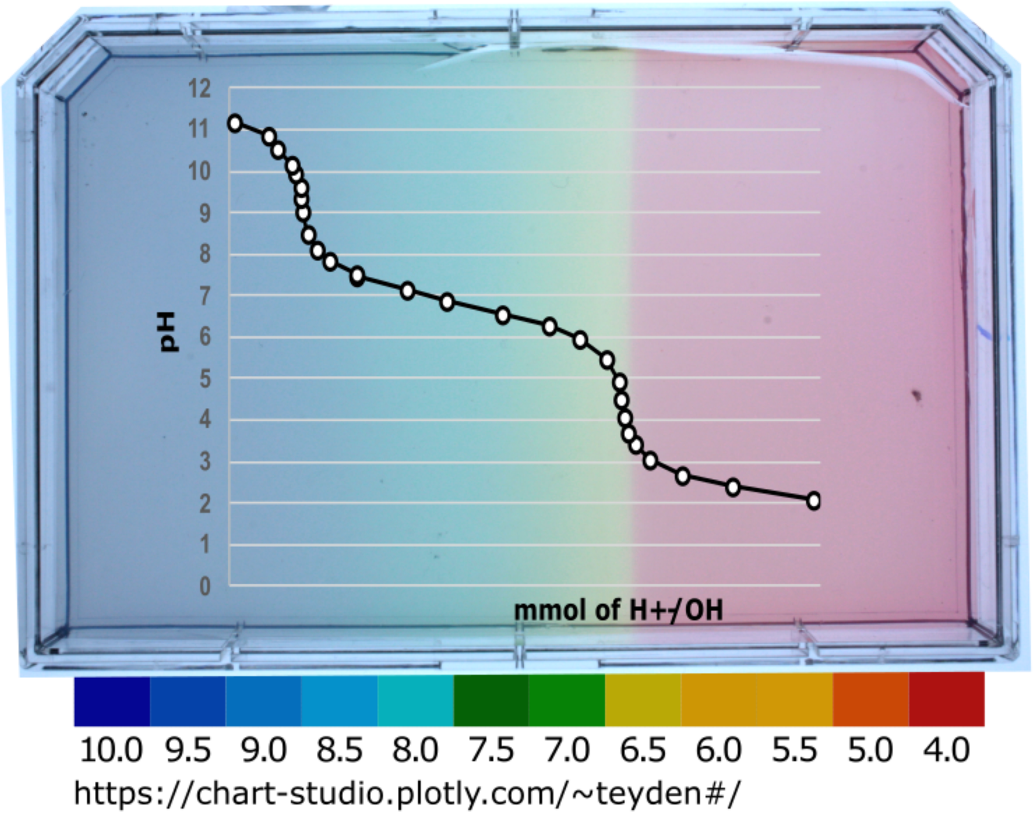


**Supplementary Figure 2:** The pH Gradient on agar media. The figure displays an agar plate stained with a universal pH indicator to visualize the pH gradient established by the addition of 60 μL of 3.3M $H_3PO_4$ to the right side and 120 μL of 10 M KOH to the left side, mirroring the conditions applied in the experiments. The plate includes a superimposed graph of a titration curve of a phosphate buffer, the main buffering agent on the media, providing a reference for interpreting the pH-dependent growth patterns observed in the *Chlamydomonas* strains.

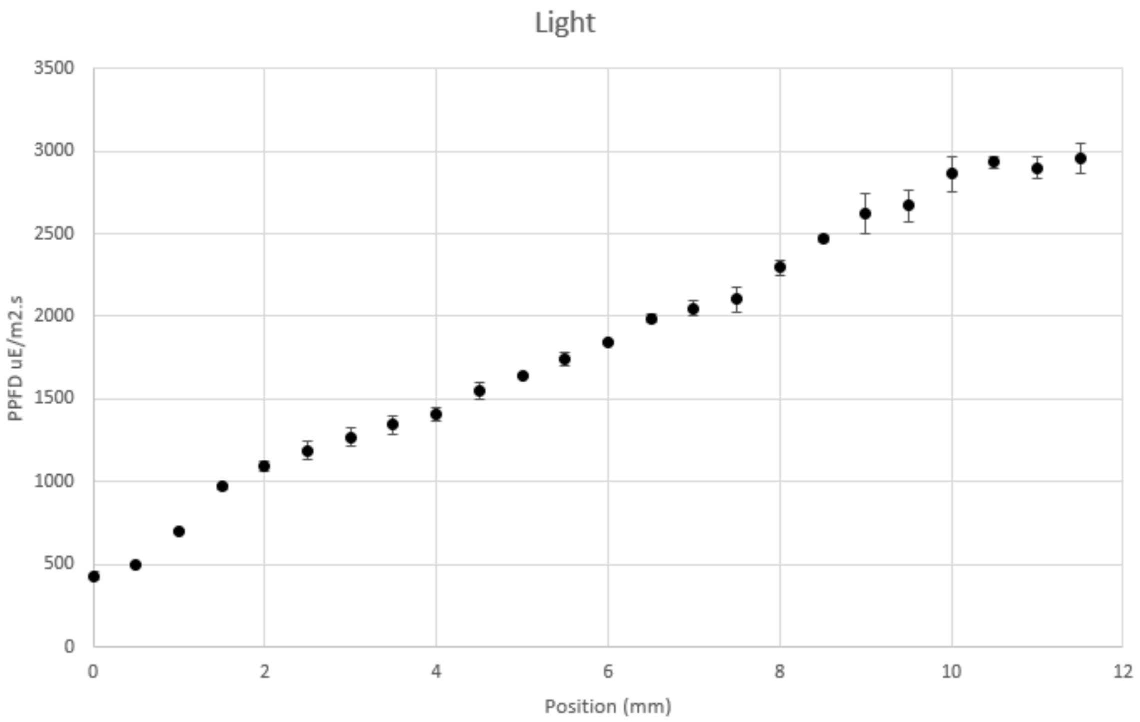


**Supplementary. Figure 3:** Light gradient the cells were exposed in HSM agar plate. Light is provided by blue and red LEDs in a 1000W Mastiff GrowL® LED grow light. PPFD was read in triplicates (n = 3) throughout the plate every 5 mm. The average results are displayed in the graph. The error bars indicate the standard deviation. The initial point was the edge of the OmniTray™ Single-Well Plate.

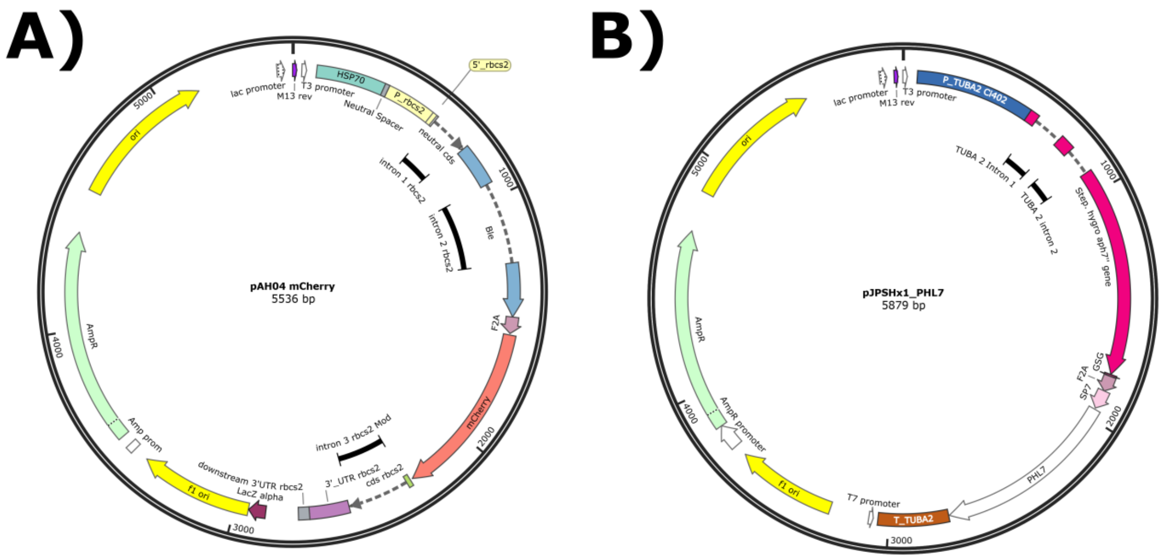


**Supplementary Figure 4**: Vectors map from **A**) The vector map of the pAH04mCherry plasmid illustrates the construct enabling cytosolic expression of mCherry in *C. pacifica 402*

(CC-5697), driven by the $P_{AR1}$ promoter and $T_{rbcS2}$ terminator from *C. reinhardtii*, and including a Bleomycin resistance gene (*Ble*) and the FMDV-2A self-cleaving peptide sequence and **B**) The vector map of the pJPSHx1PHL7 plasmid shows the construct for the secretion of PHL7 in *C. pacifica 403* (CC-5699), under the control of the $P_{Tub2}$ promoter and $T_{Tub2}$ terminator from *C. pacifica*, including a Hygromycin resistance gene (*Hyg*), a signal peptide (SP) from the SAD1p gene of *C. reinhardtii*, and the FMDV-2A. The vector has the coding sequence for PHL7.

### Supplementary Tables

**Supplementary Table 1:** Codon Usage table in *Chlamydomonas pacifica*.

| #Cod | AA | Fraction | Freq. | Number | #Cod | AA | Fraction | Freq. | Number |
|---|---|---|---|---|---|---|---|---|---|
| GCA | A | 0.121 | 15.328 | 100534 | CCA | P | 0.105 | 6.331 | 41525 |
| GCC | A | 0.376 | 47.501 | 311550 | CCC | P | 0.437 | 26.374 | 172981 |
| GCG | A | 0.395 | 49.876 | 327127 | CCG | P | 0.349 | 21.079 | 138251 |
| GCT | A | 0.108 | 13.649 | 89520 | CCT | P | 0.109 | 6.563 | 43048 |
| TGC | C | 0.896 | 15.272 | 100164 | CAA | Q | 0.107 | 4.516 | 29618 |
| TGT | C | 0.104 | 1.777 | 11654 | CAG | Q | 0.893 | 37.551 | 246285 |
| GAC | D | 0.871 | 42.882 | 281250 | AGA | R | 0.018 | 1.191 | 7814 |
| GAT | D | 0.129 | 6.35 | 41650 | AGG | R | 0.127 | 8.333 | 54654 |
| GAA | E | 0.081 | 4.449 | 29181 | CGA | R | 0.041 | 2.716 | 17811 |
| GAG | E | 0.919 | 50.768 | 332975 | CGC | R | 0.444 | 29.227 | 191690 |
| TTC | F | 0.829 | 22.867 | 149979 | CGG | R | 0.291 | 19.161 | 125673 |
| TTT | F | 0.171 | 4.724 | 30986 | CGT | R | 0.079 | 5.215 | 34203 |
| GGA | G | 0.073 | 6.502 | 42642 | AGC | S | 0.373 | 23.875 | 156593 |
| GGC | G | 0.58 | 51.779 | 339607 | AGT | S | 0.037 | 2.355 | 15449 |
| GGG | G | 0.259 | 23.112 | 151583 | TCA | S | 0.055 | 3.512 | 23035 |
| GGT | G | 0.088 | 7.839 | 51411 | TCC | S | 0.27 | 17.271 | 113273 |
| CAC | H | 0.884 | 19.938 | 130766 | TCG | S | 0.206 | 13.166 | 86355 |
| CAT | H | 0.116 | 2.61 | 17117 | TCT | S | 0.059 | 3.754 | 24619 |
| ATA | I | 0.088 | 2.382 | 15620 | ACA | T | 0.113 | 5.666 | 37163 |
| ATC | I | 0.743 | 20.054 | 131530 | ACC | T | 0.428 | 21.431 | 140561 |
| ATT | I | 0.169 | 4.55 | 29845 | ACG | T | 0.388 | 19.429 | 127429 |
| AAA | K | 0.251 | 10.425 | 68373 | ACT | T | 0.071 | 3.567 | 23398 |
| AAG | K | 0.749 | 31.106 | 204019 | GTA | V | 0.047 | 3.38 | 22169 |
| CTA | L | 0.036 | 3.676 | 24113 | GTC | V | 0.205 | 14.847 | 97375 |
| CTC | L | 0.155 | 15.648 | 102631 | GTG | V | 0.675 | 48.82 | 320201 |
| CTG | L | 0.698 | 70.385 | 461636 | GTT | V | 0.073 | 5.307 | 34807 |
| CTT | L | 0.051 | 5.149 | 33770 | TGG | W | 1 | 14.69 | 96345 |
| TTA | L | 0.007 | 0.741 | 4863 | TAC | Y | 0.917 | 22.294 | 146222 |
| TTG | L | 0.052 | 5.276 | 34601 | TAT | Y | 0.083 | 2.007 | 13161 |
| ATG | M | 1 | 22.549 | 147895 | TAA | * | 0.225 | 0.586 | 3843 |
| AAC | N | 0.909 | 22.348 | 146575 | TAG | * | 0.276 | 0.72 | 4722 |
| AAT | N | 0.091 | 2.251 | 14761 | TGA | * | 0.499 | 1.302 | 8540 |

**Supplementary Table 2:** Mating genes mapping between *C. reinhardtii* and *C. pacifica*

| Gene | Gene ID (*C. reinhardtii*) | Gene ID (*C. pacifica 402*) | E-value | Query Cover(%) | identity(%) |
|---|---|---|---|---|---|
| mtd1 | Cre06.g800601_4532.1 | anno1.g563.t1 | 5.67E-94 | 69.14446 | 31.41655 |
| mid | Cre06.g800600_4532.1 | anno1.g11755.t1 | 1.65E-09 | 51.351351 | 22.297297 |
| fus1 | Cre06.g252750_4532.1 | anno2.g10822.t1 | 1.32E-44 | 30.825243 | 11.043689 |
| sag1 | Cre08.g374250_4532.1 | anno1.g15435.t1 | 4.17E-78 | 11.1699 | 4.644327 |
| sad1 | Cre12.g517300_4532.1 | anno1.g7887.t1 | 1.79E-69 | 8.966461 | 7.734428 |
| hap2 | Cre16.g674852_4532.1 | anno1.g6770.t1 | 0.00E+00 | 55.390009 | 37.072743 |
| gsm1 | Cre08.g375400_4532.1 | anno1.g15725.t1 | 1.43E-33 | 11.44385 | 5.775401 |
| gsp1 | Cre02.g109650_4532.1 | anno2.g5263.t1 | 1.33E-41 | 21.129503 | 9.055501 |
| gex1 | Cre06.g280600_4532.1 | anno1.g2932.t1 | 3.54E-88 | 56.681351 | 25.991189 |
| mmp1 | Cre17.g718500_4532.1 | anno2.g5240.t1 | 1.17E-48 | 74.334898 | 23.943662 |

**Supplementary Table 3a**: HSM media composition

| Compound | Final mM |
|---|---|
| NH4Cl | 9.3474 |
| MgSO4 . 7H2O | 0.0811 |
| CaCl2 . 2H2O | 0.0680 |
| K2HPO4 | 8.2664 |
| KH2PO4 | 5.2908 |
| Trace Elements | 1mL/L |
| Acetic Acid | 15.3454 |
| *pH adjusted with NaOH 10M | |

**Supplementary Table 3b**: Tris-Acetate-Phosphate(TAP) media composition

| Compound | Final mM |
|---|---|
| $NH_4Cl$ | 9.3474 |
| $MgSO_4.7H_2O$ | 0.0811 |
| $CaCl_2.2H_2O$ | 0.068 |
| $K_2HPO_4$ | 0.9989 |
| $KH_2PO_4$ | 0 |
| Tris HCl | 19.9775 |
| Trace Elements | 1mL/L |
| Acetic Acid | 15.3454 |
| *pH adjusted with NaOH 10M | |

**Supplementary Table 3c**: D2-15 media composition

| Compound | Final mM |
|---|---|
| $NaNO_3$ | 8.2400 |
| $MgSO_4.7H_2O$ | 0.3000 |
| $CaCl_2.2H_2O$ | 0.1800 |
| $K_2HPO_4$ | 0.5600 |
| $KH_2PO_4$ | 0.2100 |
| $Na_2CO_3$ | 141.5 |
| Trace Elements | 1mL/L |

**Supplementary Table 3d**: **Kropat** media composition

| Compound | Final µM |
|---|---|
| EDTA-$Na_2$ | 55.5 |
| $Na_2CO_3$ | 22 |
| $FeCl_2$ | 20 |
| $MnCl_2$ | 6 |
| $ZnSO_4$ | 2.5 |
| $CuCl_2$ | 2 |
| $Na_2SeO_3$ | 0.1 |
| $(NH_4)_6Mo_7O_{24}$ | 0.0285 |

### Supplementary Videos

**Supplementary Video 1**: Motility tracking of *C. pacifica* - (Raw_402_Video_1.mp4)
**10.5281/zenodo.14059999**
**Supplementary Video 2**: Motility tracking of *C. pacifica* - (Raw_402_Video_2.mp4)
**10.5281/zenodo.14059999**
**Supplementary Video 3**: Motility tracking of *C. pacifica* - (Raw_402_Video_3.mp4)
**10.5281/zenodo.14059999**
**Supplementary Video 4**: Motility tracking of *C. pacifica* - (TrackMate capture of Video_1_402.avi)
**10.5281/zenodo.14059999**
**Supplementary Video 5**: Motility tracking of *C. pacifica* - (TrackMate capture of Video_2_402.avi)
**10.5281/zenodo.14059999**
**Supplementary Video 6**: Motility tracking of *C. pacifica* - (TrackMate capture of Video_3_402.avi)
**10.5281/zenodo.14059999**
**Supplementary Video 7**: Mating video 1 - (Starting to aggregate 402 403.avi)
**10.5281/zenodo.14059999**
**Supplementary Video 8**: Mating video 2 - (Fusing_cell_wall_motility.avi)
**10.5281/zenodo.14059999**

### Supplementary Method

**Protocol from UCR metabolomics core, UCRiverside.**

**Methods for LC-MS and GC-MS analysis of Algae Culture Supernatant and Pellet.**

##### Supernatant sample preparation

Supernatant collected from algae cultures (25 ml) was freeze-dried and resuspended in a smaller volume (10 ml) with the addition of ribitol (TCIA0171) as an internal standard (final concentration of 1.67 ug/ml). Aliquots of resuspended supernatant (0.6 ml) were added to Eppendorf tubes and freeze-dried completely.

##### Pellet sample preparation

Lyophilized pellets (9.5-10 mg) were weighed into glass tubes (VWR46610-724) and submitted to biphasic metabolite extraction.

**Biphasic metabolite extractions for Lipidomics**

Biphasic extractions were conducted, as reported in Hollin et al. (2022), with minor modifications. In short, ice-cold extraction buffer (3:2 methyl tert-butyl ether:80% methanol) was added proportionally to dry pellet (1ml/10mg) and to freeze-dried supernatant (1ml/0.6 ml of freeze-dried supernatant). Samples were sonicated for 5 min on ice, followed by 1h vortexing at 4 °C. LC-MS grade water (270 ul water / 1 ml extraction buffer) was added to induce phase separation. The water used for the pellet extraction phase separation contained ribitol as an internal standard for the GC-MS analysis (final concentration in water of 3.7 ug/ml). Samples were sonicated on ice for 5 min, vortexed briefly, sonicated again for 5 min, and vortexed for 10 min at 4 °C. Samples were centrifuged for 30 min at 1399 *g* and 4 °C. The top organic layer was collected (200 ul) for Lipidomics analysis (LC-MS), dried down under a nitrogen stream, and resuspended in 100 µL of 9:1 methanol:toluene solution for injection. The aqueous bottom layer was collected (200 ul) and dried down under a nitrogen stream for chemical derivatization and GC-MS metabolomics.

As reported previously, the dried-down aqueous phase was derivatized with minor modifications by adding 50 ul of Methoxyamine reagent (20mg/ml in pyridine) and incubating for 120 mins at 37 °C. After the first incubation, 50 ul of MSTFA +1% TMCS solution (ThermoTS48915) was added and incubated for 30 mins at 37 °C. Samples were injected (2 ul) directly into the GC-MS (Bhatia et al. 2019).

**LC-MS lipidomic analysis**

LC-MS lipidomics analysis was performed at the UC Riverside Metabolomics Core Facility. Briefly, analysis was performed on a G2-XS quadrupole time-of-flight mass spectrometer (Waters) coupled to an I-class UPLC system (Waters). Separations were carried out on a CSH C18 column (2.1 x 100 mm, 1.7 µM) (Waters). The mobile phases were (A) 60:40 acetonitrile:water with 10 mM ammonium formate and 0.1% formic acid and (B) 90:10 isopropanol:acetonitrile with 10 mM ammonium formate and 0.1% formic acid. The flow rate was 400 µL/min, and the column was held at 65 °C. The injection volume was 1 µL. The gradient was as follows: 0 min, 15% B; 2 min, 30% B; 3 min, 50% B; 10 min, 55% B; 14 min, 80% B, 16 min, 100% B; 20 min, 100% B; 20.5 min, 15% B; 30 min, 15% B.

The MS was scan range was (50 to 1600 m/z) with a 100 ms scan time. MS/MS was acquired in data-dependent fashion. Source and desolvation temperatures were 150 °C and

600 °C, respectively. Desolvation gas was set to 850 L/hr and cone gas to 50 L/hr. All gasses were nitrogen except the collision gas, which was argon. Capillary voltage was 1 kV in positive ion mode and 2 kV in negative ion mode.

A quality control sample, generated by pooling equal aliquots of each sample, was analyzed periodically to monitor system stability and performance. Samples were analyzed in random order. Leucine enkephalin was infused and used for mass correction.

**Lipidomic data analysis**

**Lipidomics**

Data analysis (peak picking, alignment, deconvolution, integration, normalization, and spectral matching) was performed on the Progenesis Qi software (Nonlinear Dynamics, Durham, NC). Data was normalized to total ion chromatogram, and features with a coefficient of variation more significant than 30% were removed. RAMClust (Broeckling et al. 2014) was used to aid in identifying features belonging to the same metabolite (i.e., in source fragments). A slightly modified version of the metabolomics standard initiative guidelines was used to assign annotation level confidence (Sumner et al. 2007; Schymanski et al. 2014): Annotation level 2a indicates a match of MS and MS/MS to an external database. Level 2b indicates a match of MS and MS/MS to in silico databases (Blaženović et al. 2019). Several mass spectral metabolite databases, including Metlin (Guijas et al. 2018), Lipidblast (Kind et al. 2013; Blaženović et al. 2019), Lipidmaps (Schmelzer et al. 2007) and HMDB (Wishart et al. 2022) were used for compound identification.
